# An unconventional cerebrospinal fluid-derived Semaphorin-signalling regulates apical progenitor dynamics in the developing neocortex

**DOI:** 10.1101/2020.05.20.106526

**Authors:** Katrin Gerstmann, Karine Kindbeiter, Ludovic Telley, Muriel Bozon, Camille Charoy, Denis Jabaudon, Frédéric Moret, Valerie Castellani

**Affiliations:** Institut NeuroMyoGène, CNRS UMR 5310, INSERM U1217, University Claude Bernard, 69008 Lyon, France; Department of Basic Neuroscience, University of Geneva, 1211 Geneva 4, Switzerland; UCL Institute of Ophthalmology, University College London, London, UK

**Keywords:** extrinsic Semaphorin/Neuropilin-complexes, CSF, apical adhesion, nuclear positioning, corticogenesis

## Abstract

In the embryonic brain, dynamic regulation of apical adhesion is fundamental to generate correct numbers and identity of precursors and neurons. Radial glial cells (RGC) in the cerebral cortex are tightly attached to adjacent neighbours. However, cells committed to differentiate reduce their adhesiveness to detach and settle at distal position from the apical border. Whether diffusible signals delivered from the cerebrospinal fluid (CSF) contribute to the regulation of apical adhesion dynamics remain fully unknown. Here we report that unconventional pre-formed complexes of class3-Semaphorins (Sema) and Neuropilins (Nrp) are released into the cerebrospinal fluid (CSF) from sources including the choroid plexus. Through analysis of mutant mouse models and various ex vivo assays, we propose that two different complexes, Sema3B/Nrp2 and Sema3F/Nrp1, bind to apical endfeet of RGCs, and exert dual regulation of their attachment, nuclei dynamics, that oppositely promotes or inhibits basal progenitor and neuron differentiation. This reveals unexpected contributions of CSF-delivered guidance molecules during cortical development.

## Introduction

During development of the vertebrate central nervous system, cell cycle kinetics and cell fate decisions of neuronal progenitors are precisely orchestrated by complex intrinsic and extrinsic mechanisms. Cortical stem cells are located in the ventricular zone and exhibit a highly polarized morphology with a basal and an apical process that are anchored to the pial basement membrane and the ventricular surface, respectively ^1^. The apical endfeet are in direct contact with the cerebrospinal fluid (CSF), allowing cortical stem cells to receive extrinsic signals from the cerebral ventricles ^2^. During early corticogenesis, apical progenitors divide symmetrically to expand the pool of progenitor cells ^3,4^. However, at the onset of neurogenesis they divide asymmetrically to generate both proliferative and postmitotic progeny. Apical endfeet are tightly attached to adjacent neighbours via adherent junctions and cells that are committed to differentiate reduce the apical connections to disengage from the apical surface ^5–7^. The detachment is essential for further neurogenesis and migration of the differentiated cells ^8,9^. Apical progenitors are highly dynamic along the apico-basal axis and undergo interkinetic nuclear migration (INM) in synchrony with their cell cycle. The nucleus oscillates from the apical pole where mitosis occurs, to a more basal position where S-phase is achieved. Apical endfeet, nuclear dynamic and cell cycle kinetics are considered as major parameters determining the balance between proliferation and neurogenesis are crucial for cortical integrity ^10^. Nevertheless, the developmental mechanisms and factors controlling the spatial architecture of apical progenitors and their dynamic are vastly unknown. The apical endfeet are in direct contact with the cerebrospinal fluid (CSF) of the ventricles, from where they receive extrinsic signals ^2^. Recently, it has been discovered that proteins acting as guidance cues for migrating cells and axons, also influence neural progenitor proliferation and differentiation ^11–13^. Class 3 Semaphorins are secreted proteins exerting either repulsive or attractive effects upon binding to transmembrane receptor complexes composed of Neuropilins and Plexins ^14^. We recently observed that the Semaphorin3B (Sema3B) is released into the CSF by the nascent choroid plexus and the floorplate, where it influences the proliferation and division orientation of progenitor cells in the developing spinal cord ^15^. This broaches the issue whether soluble Semaphorins in the CSF influence cortical progenitor cells as well.

Here we provide evidence for an unconventional Sema/Nrp-signalling pathway that controls the nuclear positioning and cell adhesion of cortical progenitor cells. Our data suggest that Class3-Semas and soluble Nrps are expressed by the embryonic choroid plexus and released into the cerebrospinal fluid. They form specific complexes that bind to Plexins, which are present on apical progenitor cells. The resulting signalling exerts collaborative efforts in regulating the apical positioning of mitotic nuclei at the ventricular border and the adhesive properties of these cells. Our results indicate that Sema3B/Nrp2-signalling causes pro-apical forces and increases adhesion, that reduces the generation of differentiated cells. In turn, Sema3F/Nrp1-signalling exerts anti-apical forces and decreases adhesion needed for setting transient amplifying cells and postmitotic neurons. Altogether our results suggest that extrinsic CSF-derived Sema/Nrp-complexes are crucial for apical dynamic of neural progenitors and contribute to the proper generation of subsequently differentiated cells in the developing cerebral cortex.

## Results

### Class3-Semas are expressed by the embryonic choroid plexus and released into the CSF

To reveal the expression of Sema3s in the developing cerebral cortex, we performed *in situ* hybridisations on brain sections at different embryonic stages, focusing on Sema3B and Sema3F. We detected both transcripts in the nascent choroid plexus of the lateral ventricles starting from E11.5, when the formation of this structure is initiated ^16^, with abundant expression of Sema3B and Sema3F mRNA reached at E13.5 (Figure 1A). Notably, we observed no signal in the cortical wall. In addition, we detected Sema3B and Sema3F transcripts in the developing choroid plexus of the 4^th^ ventricle starting from E10.5 (Figure 1B). The data suggest that Sema3B and 3F are expressed by the developing choroid plexus, but not by cortical progenitor cells in the ventricular zone of the telencephalon. These findings were confirmed by real-time functional single-cell transcriptome analysis of cortical cells according to Telley et al., (2016), in which no Class3-Semaphorins were detected in apical progenitors (Figure 1E,F). Since no suitable antibodies are available for immunostaining of Sema3B and Sema3F, we took advantage of a Sema3B-GFP knock-in (ki) mouse line ^15^ to reveal the endogenous Sema3B-expression. In this mouse model, the N-terminus of Sema3B is fused to an enhanced green fluorescent protein (GFP), which can be detected with an anti-GFP antibody. GFP-immunolabeling confirmed the prominent expression of endogenous Sema3B-GFP in the nascent choroid plexus of Sema3B-GFP ki embryos at E12.5 (Figure S1A) and E13.5 (Figure 1C). Moreover, Sema3B-GFP appeared to be concentrated at the apical border of the CP, suggesting that Sema3B is secreted into the CSF. To confirm the presence of Class3-Sema molecules in the cerebrospinal fluid, we performed anti-GFP western blot experiments on CSF samples harvested from the embryonic brain ventricles at E13.5 of Sema3B-GFP ki or wildtype embryos as control. We indeed detected Sema3B-GFP in the CSF of Sema3B-GFP ki mice (Figure 1D). Altogether our results suggest that the nascent choroid plexus is a main source for Sema3B and Sema3F during cortical development and releases these molecules into the CSF. We thus examined whether cortical progenitor cells express Neuropilins and Plexins, which form receptor complexes mediating the Sema3-signalling.

**Figure 1:**
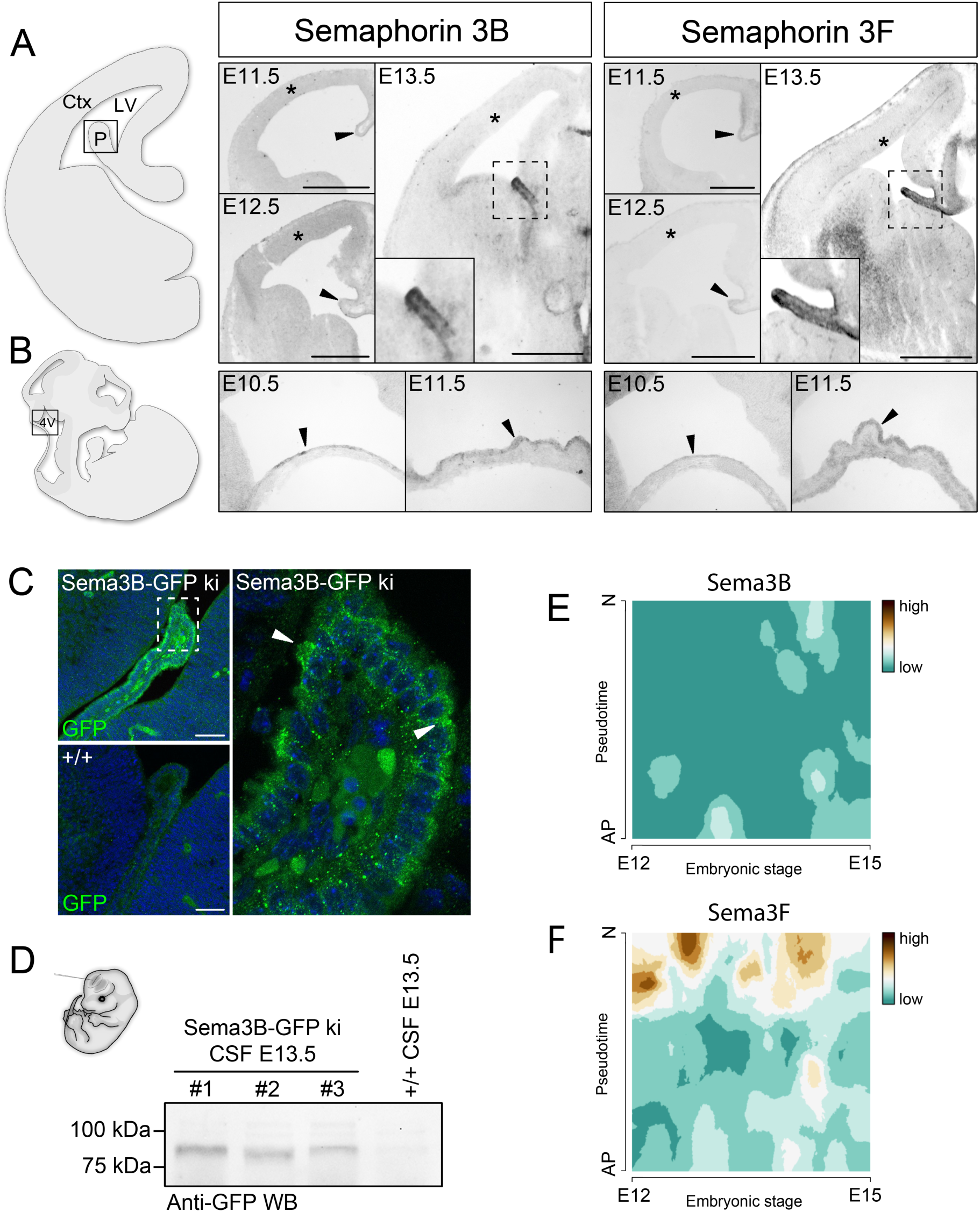
Class3-Semaphorins are expressed by the developing choroid plexus and secreted into the CSF. **(A, B)** *In situ* hybridisations on coronal sections through the telencephalon and sagittal sections of the midbrain show the mRNA expression of Sema3B and Sema3F at indicated ages. Arrows point to the developing choroid plexus. Asterisks indicate the cortical ventricular zone, which is composed of apical progenitors. **(C)** Fluorescence micrographs showing GFP-labelling of the nascent choroid plexus in Sema3B-GFP ki/ki and WT embryos at E13.5. Arrows point to the GFP-accumulation at the apical border of the choroid plexus. **(D)** GFP-immunoblotting on CSF from Sema3B-GFP ki/ki embryos shows the presence of soluble Sema3B-GFP molecules in the brain fluid. **(E, F)** Single cell transcriptome sequencing of embryonic cortical cells shows that neither Sema3B (**E**), nor Sema3F **(F)** are expressed by cortical progenitor cells. Pseudotime describes the differentiation from an apical progenitor (0) to a postmitotic neuron (1) between E12 and E15. Scale bars: 500 µm **a** and 100 µm **b**. Ctx, Cortex; LV, lateral ventricle; P, choroid plexus.

### Soluble Neuropilins are released in the CSF and form complexes with Sema3s, that bind to Plexins at apical progenitors

We first investigated the expression of the Neuropilin receptors, Nrp1 and Nrp2, using *in situ* hybridisation and immunolabelling (Figure 2A) on embryonic brain sections at E13.5. With both approaches, we observed Nrp1 and Nrp2 in the cortical plate of the dorsal telencephalon, which reveals Nrp expression in differentiating neurons. Strikingly however, we detected neither Nrp1 nor Nrp2 mRNA in the cortical ventricular zone, where the proliferating progenitor cells are located (Figure 2A). These findings were confirmed by real-time functional single-cell transcriptome analysis, indicating that unlike differentiating cortical cells, apical progenitor cells express none of these receptor sub-units (Figure 2B,C).

**Figure 2:**
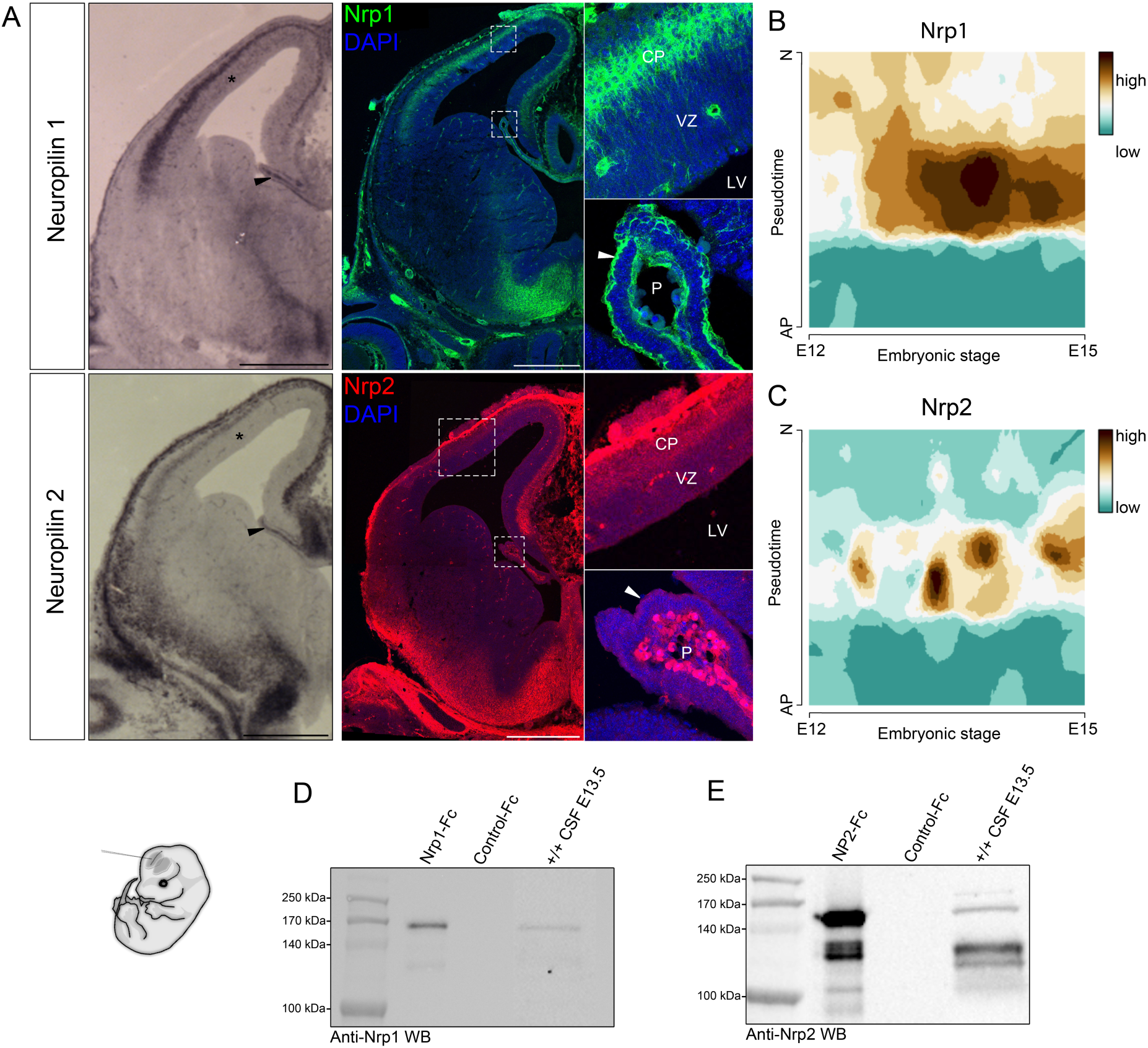
Neuropilins are expressed by the developing choroid plexus and released in the CSF. **(A)** *In situ* hybridisation and immunolabelling of Nrp1 and Nrp2 on embryonic brain sections at E13.5 show the expression of the Class3-Semaphorin receptors in the nascent choroid plexus. No signal was detected in the cortical ventricular zone. Black arrowheads point to the developing choroid plexus. The white arrowheads point to Nrp1/Nrp2 protein stainings at the apical border of epithelial cells in the developing choroid plexus. Asterisks indicate the cortical progenitor regions. **(B, C)** Single cell transcriptome analysis of embryonic cortical cells shows that cortical progenitor cells express neither Nrp1 **(B)**, nor Nrp2 **(C)**. Pseudotime describes the differentiation from an apical progenitor cell (0) to a postmitotic neuron (1) between E12 and E15. **(D, E)** Immunoblotting on embryonic CSF using antibodies against Nrp1 (**D**) and Nrp2 (**E**) indicates that both receptors are released into the brain fluid. Recombinant Nrp1 and Nrp2 ectodomains fused to Fc fragment and are used as positive controls of immunodetection. Scale bars, 500 µm. CP, cortical plate; LV, lateral ventricle; VZ, ventricular zone.

Interestingly, in situ hybridization revealed the expression of both Nrp1 and Nrp2 in the developing choroid plexus of the lateral ventricle (Figure 2A). Close examination of Nrp1 staining (Figure 2A) indicates the presence of Nrp1 in vessel wall cells located in the stroma of these choroid plexi, which is reminiscent of expression of Nrp1 in endothelial reported in other systems ^18^. Intriguingly, anti-Npr1 immunostaining detected Nrp1 proteins not only in the stroma of the telencephalic choroid plexi but also at the apical borders of their epithelial cells. In situ hybridization also revealed Nrp2 mRNA in the stromal compartment of the telencephalic CP, suggesting its expression in vessel walls. We then compared Nrp2 protein expression by immunolabeling of E13.5 Nrp2+/+ and Nrp2-/- brains and detected Nrp2 at the choroid plexus apical border. (Figure 2A, S1C). In addition, we also observed strong Nrp2 expression, both at the transcript and protein levels, in the meninges surrounding the cortex and in cell populations in contact with ventricles in several regions of the diencephalon and the midbrain, particularly the floor plate (Figure S1C-D). The specificity of Nrp2 labelling was validated by immunolabeling performed on Nrp2-/- brains (Figure S1C). For Nrp1, we had available Nrp1^Sema/Sema^ line that does not allow such validation since the protein lacks Sema3 binding but is still present. Nevertheless, immunolabeling with the secondary antibody alone do not reveal an apical staining of the choroid plexus (Figure S1B). These observations raised the possibility that Nrps, instead of being expressed by the progenitors themselves, would be released as soluble forms, as their ligands. For both Nrps, soluble secreted forms generated by ectodomain shedding and encoded by specific splicing variants have previously been reported ^19,20^. To address whether soluble Nrps are released from the choroid plexi, meninges and other sources of the neural tube wall, we thought to explore their presence in the CSF. We performed western blot analysis of the CSF harvested from embryonic brain ventricles of E13.5 wildtype embryos, using specific antibodies against Nrp1 and Nrp2 characterized in previous studies ^14,21^. We detected several forms for both Nrp1 (Figure 2D) and Nrp2 (Figure 2E) in the CSF with molecular weights consistent with Nrp ectodomains. Thus, both Sema3s and Nrps are released into the embryonic brain ventricles and are available for binding to cortical progenitor cells. We wondered whether pre-formed complexes are present in the CSF. Because anti-Sema3B or anti-Sema3F antibodies were not specific enough to detect small amount of proteins from *in vivo* samples, co-immunoprecipitation experiments were performed with CSF from embryonic brains of conditional Sema3B-GFP ki and WT embryos at E13.5. Immunoprecipitation was performed using anti-GFP antibodies and followed by western blot against Nrp1 or Nrp2 (Figure 3A). The results revealed Nrp2/Sema3B-complexes in the CSF of Sema3B-GFP ki embryos. The sizes of Nrp2 isoforms coimmunoprecipitated with Sema3B are similar to Nrp2 ectodomains of the recombinant Nrp2-Fc control. A weak signal was also detected for the Nrp1/Sema3B-condition, suggesting that Nrp1/Sema3B-complexes if present might be much rare.

**Figure 3:**
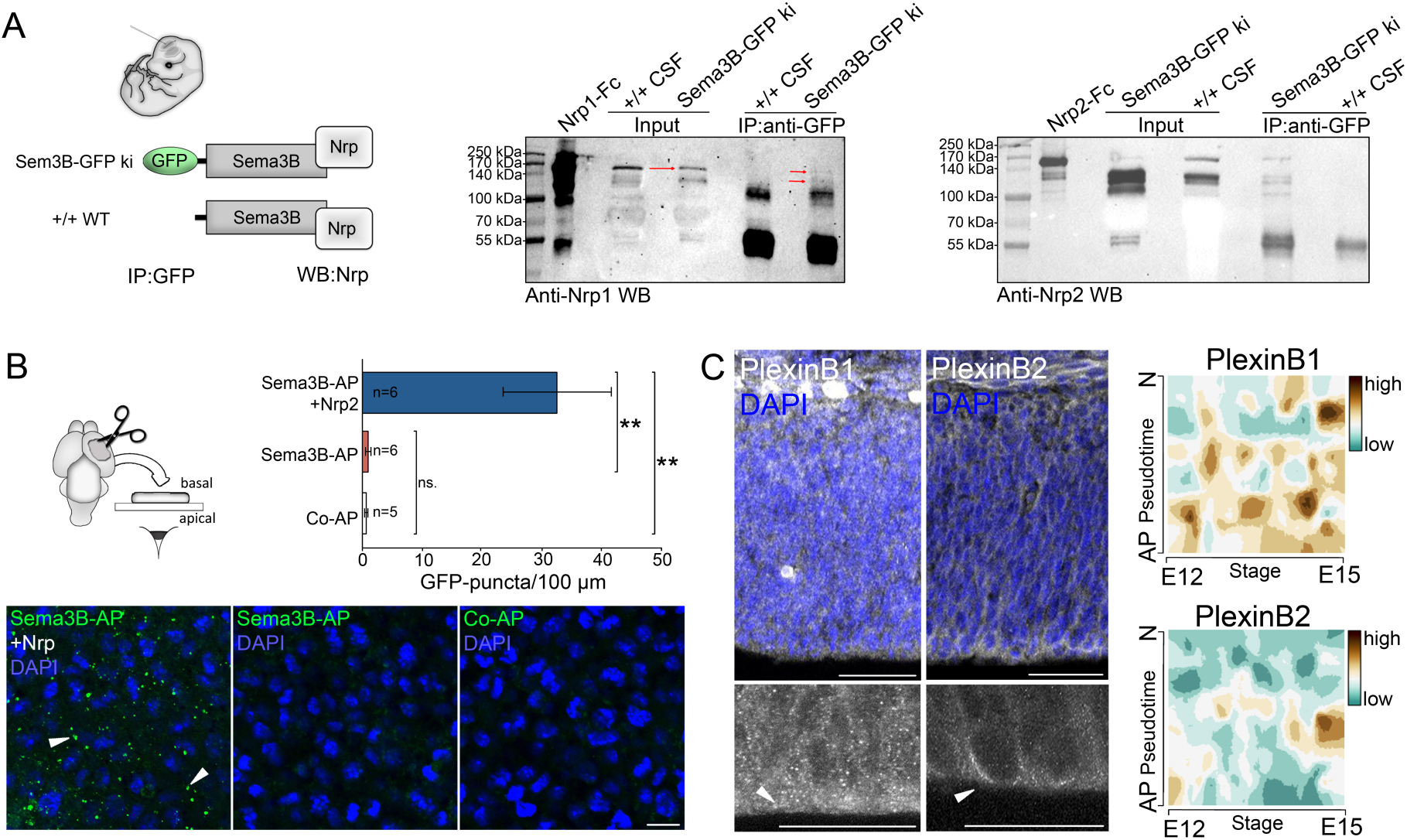
Sema3s and Neuropilins form complexes in the embryonic CSF that bind to Plexins at the apical endfeet of radial glia cells. **(A)** Immunoprecipitation of GFP using CSF of Sema3B-GFP ki/ki mice and WT embryos show prominent binding of Sema3B-GFP with Nrp2, but rare association with Nrp1 in the CSF. **(B)** *En face* binding assays reveal that exposure of cortical tissue to recombinant Sema3B-AP and Nrp2-Fc results in binding of the protein complex to the apical surface. In contrast, Sema3B-AP alone and control-AP do not bind to the apical surface. The histogram depicts the quantification of AP puncta. **(C)** Immunolabeling of PlexinB1 and PlexinB2 on cortical sections at E12.5 show the expression of these Semaphorin receptors in the developing cortex. Arrows point to the accumulation of the Plexin proteins at the apical surface. Single cell transcriptome analysis of embryonic cortical cells reveals an expression of PlexinB1 and PlexinB2 in apical cortical progenitor cells. Pseudotime describes the differentiation from an apical progenitor cell (0) to a postmitotic neuron (1) between E12 and E15. Notably, no other Plexins were detected in proliferating cortical cells (Figure S1E). Mean ± SEM (n= analysed hemispheres out of two litters); t-test, ** p<0.01.

To test whether Sema/Nrp-complexes bind to the apical surface of cortical progenitor cells, we performed *en face* binding assays with recombinant proteins. For this approach, hemispheres of E13.5 wildtype embryos were separated and exposed to recombinant Sema3B fused to Alkaline Phosphatase (AP) together with combined Nrp1-Fc/Nrp2-Fc or control-Fc. After AP-immunolabelling, the dorsal telencephalon was dissected and flat mounted. Notably, images from the apical surface display the presence of punctate AP-staining exclusively in the condition where Semas were incubated together with combined Nrp1-Fc/Nrp2-Fc (Figure 3B). Neuropilin recruits signalling co-receptors among Plexin family members ^22^ that trigger intracellular signalling upon Sema3 binding. Interestingly, apical progenitor cells were reported to express PlexinB1 and PlexinB2 and the deletion of these co-receptors causes impaired cortical neurogenesis ^23^. By real-time functional single-cell transcriptome analysis, we found transcripts for PlexinB1 in apical progenitor cells (Figure 3C). Expression of other Plexins were limited if absent. Modest signal for PlexinA1 and PlexinB2 were noted (Figure 3C, Figure S1). Immunolabeling of E12.5 embryonic cortical sections against PlexinB1 and PlexinB2 confirmed the expression at protein levels in apical progenitor cells (Figure 3C). Interestingly, the results revealed their presence at the feet of cells anchored at the apical border, thus at expected location to interact with the CSF-derived Sema/Nrp-complexes (Figure 3C). Altogether these results suggest that CSF-derived Nrps enable secreted Sema3s to bind to Plexins that are present on the apical surface of cortical progenitor cells.

### CSF- derived Sema/Nrp-signalling is dispensable for self-renewal of cortical stem cells

Previous work by our group has demonstrated that Sema3B delivered in the central canal regulates the division orientation of progenitor cells in the spinal cord ^15^. Therefore, we examined whether Sema/Nrp-complexes in the embryonic brain ventricles also contribute to the division orientation of cortical progenitor cells with respect to the apico-basal axis. First of all, we examined the *in vivo* consequences of loss of Sema/Nrp-signalling in Sema3B ko ^14^, Sema3F ko ^24^, Nrp2 ko ^25^, and Nrp1^Sema/Sema^ mice, in which the Sema3 binding is selectively disrupted ^26^, respectively. Coronal embryonic brain sections of E12.5 embryos were labelled with antibodies against phospho-histone 3 (PH3), to stain cells undergoing mitosis, and against γTubulin to label the spindle poles (according to Arbeille et al., 2015). Altogether our investigations revealed no significant changes in the division orientation of apical progenitor cells in the developing cerebral cortex of the analysed transgenic mouse lines (Figure S2). These results indicate that the CSF-derived proteins do not play a crucial role for the division orientation of apical cortical progenitor cells in the dorsal telencephalon, at least at the examined stage and in the analysed cortical region.

To more directly assess the functional properties of Sema3B/3F on the self-renewal and multipotency of cortical progenitor cells, we also performed a classical *in vitro* proliferation experiment, the neurosphere assay. For this approach, cortical single cell suspensions from embryonic WT brains at E12.5 were prepared and subsequently exposed to different combinations of recombinant Sema-Fc and Nrp-Fc proteins, respectively. After 7 days in culture (7div) characteristic free-floating cell clusters (neurospheres) were formed by proliferating neural stem cells. The size and numbers of the resultant spheres were measured as indicators of whether the added recombinant proteins exert proliferative effects. We observed no significant differences between the conditions suggesting that the Sema/Nrp-signalling do not affect stem cell proliferation *in vitro* (data not shown). These findings were in addition confirmed by immunolabelling of embryonic brain sections at E12.5 with specific antibodies against Sox2 (sex-determining region Y-box containing gene 2) that stains apical progenitors, and PH3, a marker for mitotic cells, in Sema3B, Sema3F, Nrp1 ^Sema/Sema^ and Nrp2 mutant mice. Analysis of the quantity and distribution of Sox2-positive cells revealed no respective changes in number or distribution in these mouse mutant lines at E12.5 in comparison to wildtype littermates (Figure S3A). In addition, no differences were observed in the number of PH3-reactive nuclei in the analysed mutant embryos (Figure S3B). These findings support the conclusion that the CSF-derived Sema3B/3F-Nrp-signalling is dispensable for the expansion and renewal of the embryonic cortical apical progenitor pool.

### Sema/Nrp-interactions control the adhesive properties of apical progenitor cells

Nrps were initially characterized as membrane proteins mediating adhesion and several studies have highlighted interplays between Semaphorins and Neuropilins in cell adhesion in various contexts ^27–29^. Considering the accessibility of apical adhesion points of apical progenitor cells to CSF-derived cues, we thus hypothesized that the Sema/Nrp-signalling could be implicated in the regulation of progenitor apical attachment and positioning. To investigate Sema/Nrp-mediated adhesion in cortical progenitor cells *in vitro*, cortical single cells of E12.5 cortices were prepared and cultured on coverslips that were coated with recombinant Neuropilin-Fc proteins or control-Fc, in absence of other coating substrate (Figure 4A). Recombinant Semaphorin-Fc proteins or control-Fc were added to the culture medium. After 3 hours *in vitro* (3hiv), we quantified the numbers of Sox2-positive undifferentiated stem cells that adhere to Neuropilin on the coverslips. Exposure to recombinant Sema3F-Fc on Nrp1-Fc-coated coverslips resulted in significantly less Sox2-positive cells binding to the surface (Figure 4A). Conversely, Sema3B-Fc application in the Nrp2-Fc-coated condition resulted in decreased number of bound cells (Figure 4A). Thus, these data indicate that Nrp2/Sema3B-complexes can hang apical progenitors. In contrast Nrp1/Sema3F-complexes act as a non-permissive adherent substrate.

**Figure 4:**
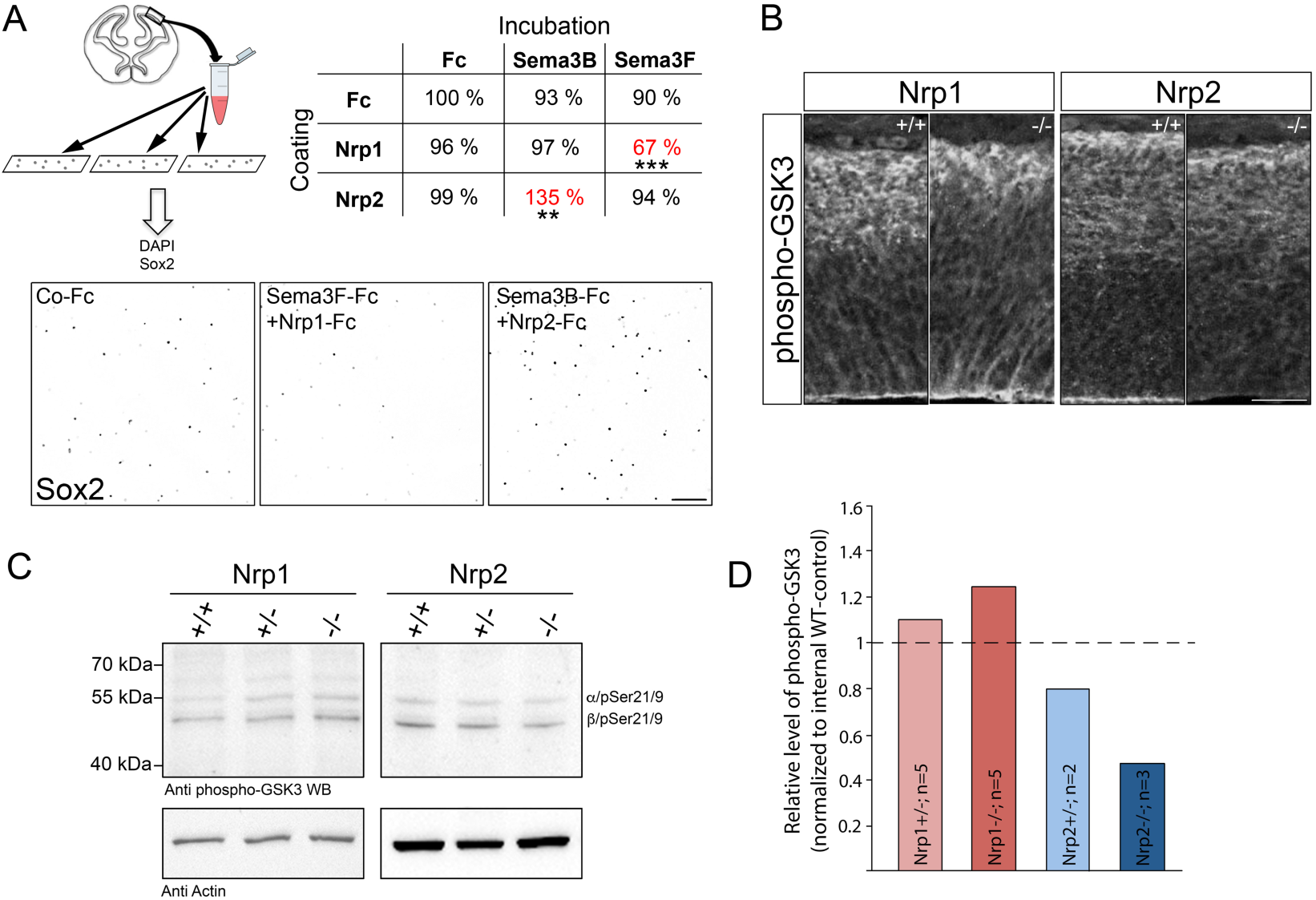
Dual effects of Semaphorin/Neuropilin-complexes on cortical progenitor adhesion and GSK3-activity in the developing cerebral cortex. **(A)** Adhesion assays on coverslips coated with recombinant Neuropilin-Fc and Semaphorin-Fc molecules in different combinations show that Sema3F-Fc/Nrp1-Fc inhibits the attachment of Sox2-positive nuclei to the substrate whereas Sema3B-Fc/Nrp2-Fc promotes cellular attachment. **(B)** Immunolabelling of inactive phospho-GSK3 forms in E12.5 embryonic brain sections prepared from Nrp1 and Nrp2 mutant embryos show an accumulation at the apical side. The immunolabeling reveals that GSK3 is inhibited in Nrp1 mutants and activated in Nrp2 knockout cortices. **(C)** Western blot of inactive phospho-GSK3 and actin on E13.5 cortical extracts of Nrp1 and Nrp2 mutant embryos show the detection of phosphorylated GSK-3α/β (Ser21/9) bands. **(D)** The quantification of the western blot signal normalized to internal wild-type controls reveals more GSK3-inhibition in Nrp1 mutants and less GSK3-inhibition in Nrp2 knockout cortices. Scale bars A: 100µm, B: 25 µm; n= number of analysed embryos.

Throughout the neurogenesis period, apical end feet undergo constant remodeling as RGCs divide, newly born cells with RGC fate re-establish apical adhesion and daughter cells committed to differentiate detach from the apical side and adjacent neighbours. In the *in vivo* context of an apical delivery, our findings suggested that both CSF-derived complexes could regulate apical anchor dynamics of cortical progenitor cells in an opposite manner. To further assess this possibility, we investigated the activity of signaling known to mediate Semas functions and to regulate adhesion. Glycogen synthase kinases (GSK3) regulate several major signaling during brain development ^30^ and thus represent appealing candidates. First, several studies reported that GSK3s act downstream of the Semas in the context of axon guidance ^31–33^. We also found in previous work that Sema3B regulates the orientation of spinal cord progenitor divisions via GSK3 signaling. Interestingly, GSK3 activity has been shown to be indispensable for proper dynamics of apical glia cell scaffold in the developing cerebral cortex ^34^. GSK3 promotes the degradation of ß-catenin in neural progenitors, a protein enriched at the apical endfeet controlling their adhesion and dynamic in coordination with N-cadherin and actomyosin complexes ^9,35^. GSK3 proteins were also proposed to be essential regulators of proliferation and differentiation during corticogenesis ^36^. These findings motivated us to analyze GSK3-activity in the cerebral cortex of Nrp mutant embryos. Immunolabelling was performed on embryonic brain sections of Nrp1 and Nrp2 mutant mice at E12.5, with anti phospho-GSK3 antibody recognizing the inactive GSK3 α and β pSer21/9 form (Figure 4B). A clear phospho-GSK3 signal was observed in the apical endfeet of cortical progenitors that appeared to be reduced in Nrp2^-/-^ cortices and augmented in the Nrp1^-/-^ brains, compared to controls. To quantify GSK3 levels, we performed western blot on cortical extracts of the different genotypes. Interestingly, this analysis confirmed that inactive forms of GSK3 were increased in Nrp1^-/-^ cortices and oppositely decreased in Nrp2 -/- brains. This also holds true in embryos lacking only one gene copy (Figure 4C-D). β-catenin mediated adhesion is negatively regulated by GSK3 activity. Thus, increased inactive GSK3 might result in increased adhesion, as expected for loss of Nrp1, while conversely, decreased inactive GSK3 might lead to increase adhesion, as expected for Nrp2. Thus, the changes of GSK3 activity observed in the Nrp mutant embryos are consistent with the dual pro- and anti-adhesive effects of Nrp2/Sema3B and Nrp1/Sema3F that we observed in our *in vitro* device.

### Sema/Nrp-signalling modulates the positioning of mitotic progenitors

It has already been described that inhibition of apical adhesion results in increased numbers of mitotic nuclei that are mislocated at distance from the ventricular border ^37,38^. We thus further investigated whether invalidation of Semas and Nrps affects the location of mitotic nuclei respective to the VZ. To approach this question, we performed an antibody labelling against PH3 in E12.5 embryonic cortical slices of the different mutant mouse lines. The overall number of PH3-positive nuclei are not altered in the absence of Sema/Nrp-molecules (Figure S3B). We defined three classes of mitotic nuclei according to their position in the cortical wall (adapted from Arvanitis et al., 2013): ventricular nuclei close to the ventricular surface, non-adjacent nuclei that were more than one cell diameter away from the ventricular border, and basal nuclei that represent intermediate cells (Figure5A).

**Figure 5:**
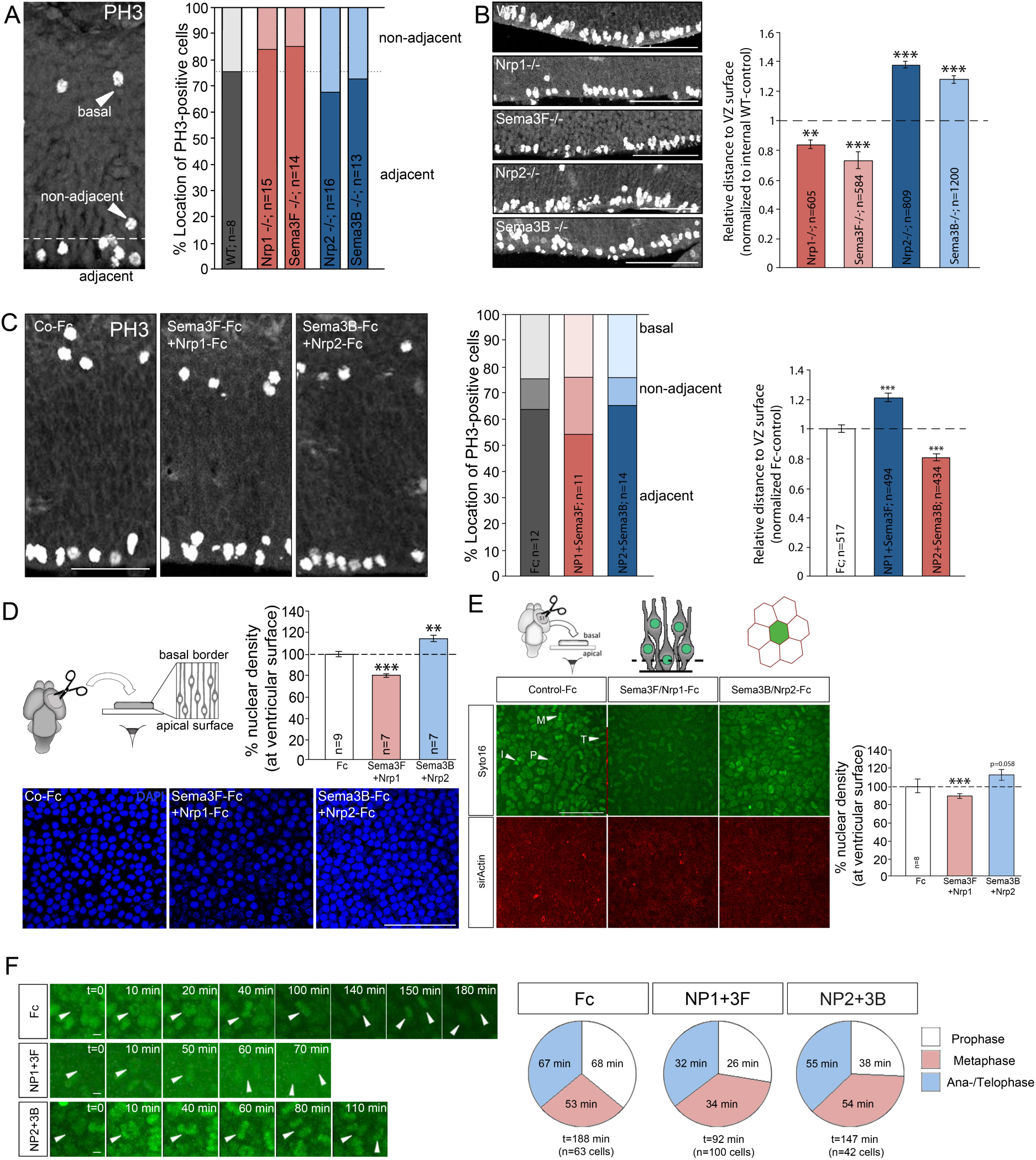
Apically-delivered Sema/Nrp-complexes regulate the positioning of mitotic nuclei and duration of mitosis. **(A)** PH3-positive nuclei were classified as ventricular, displaced or basal, according to their position in the cortical wall. The quantification reveals that Nrp1^Sema/Sema^ and Sema3F knockout mice have more ventricular nuclei and less displaced nuclei. In turn, Nrp2 and Sema3B knockout embryos have more displaced nuclei, at the expense ventricular nuclei. **(B)** Quantification of the relative distance between PH3-positive nuclei in the VZ and the ventricular surface shows that cells in Nrp1^Sema/Sema^ and Sema3F knockout embryos are closer to the ventricular surface than in control animals. In turn, mitotic nuclei in Nrp2 and Sema3B deficient embryos are more distant from the apical border than in controls. **(C)** Injections of recombinant proteins into the lateral ventricle of wildtype embryos at E13.5 regulate the positioning of mitotic cells relative to the ventricular surface. Injection of Sema3F-Fc/Nrp1-Fc results in augmented numbers of non-adjacent nuclei that are more distant to the apical surface. In turn, injections of Sema3B-Fc/Nrp2-Fc results in more adjacent PH3-positive nuclei that are closer to the ventricular surface then in control-Fc injected brains. **(D)** The positioning of apical progenitor cells can be influenced by recombinant Semaphorin-Fc and Neuropilin-Fc molecules *in vitro*. In cortices apposed to coverslips coated with Sema3F-Fc/Nrp1-Fc, less nuclei are present on the apical surface. In turn, exposure to Sema3B/Nrp2-Fc results in more nuclei present on the apical surface. **(E)** *En face* imaging of cortical tissue at E13.5 was performed to analyse the duration of mitosis of apical progenitors at the ventricular surface in the presence of recombinant Sema-Fc/Nrp-Fc proteins. The data confirm that cortical tissue apposed to Sema3F-Fc/Nrp1-Fc reveals less apical nuclei, whereas Sema3B-Fc/Nrp2-Fc exposure results in augmented numbers of apical nuclei, compared to control-Fc. **(F)** Histograms depict the absolute time of division phases of radial glia cells on the apical side of cortical tissue. An average division time of 188 min was determined under control condition. This average dropped to 92 min in the Nrp1-Fc/Sema3F-Fc condition and to 147 min in the Nrp2-Fc/Sema3B-Fc condition. The incubation with both, Sema3F-Fc/Nrp1-Fc or Sema3B-Fc/Nrp2-Fc resulted in accelerated prophase and ana-/telophase. In contrast, metaphase is accelerated upon exposure to Sema3F-Fc/Nrp1-Fc, but not to Sema3B-Fc/Nrp2-Fc. Scale bars: **A-D** 100 µm, **E** 50 µm, **F** 5 µm. T-test, * p<0.05, ** p<0.01, *** p<0.001.

The results revealed more mitotic nuclei next to the ventricular surface in Sema3F ko and Nrp1^Sema/Sema^ mice and less non-adjacent nuclei (Figure 5A). In WT embryos, we quantified 67.02 ± 2.61 % adjacent versus 21.38 ± 2.55 % non-adjacent PH3-positive cells. In contrast, Nrp1^Sema/Sema^ mice reveal 77.5 ± 1.98 % (***, p<0.001) adjacent versus 14.51 ± 1.86 % (**, p<0.01) non-adjacent and Sema3Fko mice 75.42 ± 1.23 % (p=0.14) adjacent versus 13.05 ± 1.45 % (*, p<0.05) non-adjacent nuclei. Oppositely, mitotic nuclei in Sema3B and Nrp2 knockout cortices were radially more scattered in the cortical wall than in the WT, with augmented numbers of basal PH3-positive cells. We quantified 56.19 ± 2.25 % (p=0.11) adjacent, 26.57 ± 1.92 % (p=0.59) non-adjacent nuclei in Nrp2 ko mice and 61.89 ± 2.68 % (*, p<0.05) adjacent, 22.86 ± 1.99 % (**, p<0.01) non-adjacent nuclei in Sema3B deficient embryos.

In addition, we measured the distance between mitotic nuclei in the ventricular zone and the ventricular surface (Figure 5B). In comparison to WT littermates, the mitotic nuclei in the VZ of Sema3F -/- and Nrp1^Sema/Sema^ were closer to the ventricular surface for about 22.36 ± 3.05 % (***, p<0.001) and 26.49 ± 2.78 % (***, p<0.001), respectively. In contrast, the more distant mitotic nuclei in Sema3B-/- and Nrp2 -/- cortices were reflected in an extended distance to the ventricular surface. Sema3B deficient embryos exhibited an increased distance of PH3-positive nuclei of about 28.12 ± 2.28 % (***, p<0.001) and Nrp2 deficient embryos of 38.05 ± 3.41 % (***, p<0.001). Altogether these data indicate that the position of apical progenitor nuclei during mitosis is influenced by CSF-derived signals and support that two Sema/Nrp-signalling exerts collaborative effort in balancing pro- and anti-apical forces to set nuclear position of dividing apical progenitor cells.

To confirm these functional properties of Sema/Nrp-signalling, we next created a gain-of-function approach and assessed whether Sema/Nrp-exposure affects nuclear positioning at the apical surface. We isolated E12.5 wildtype embryos and injected recombinant Sema3/Nrp proteins into their lateral ventricles. After 30 min *in vivo*, brains were fixed, sectioned, and labelled against PH3. The position of mitotic nuclei was classified as in the previous experiment (Figure 5C). Injection of control-Fc resulted in 63.92 ± 1.45 % of nuclei adjacent to the lateral surface, 11.68 ± 0.94 % of non-adjacent nuclei. Gain of Nrp1-Fc/Sema3F-Fc resulted in increased numbers of non-adjacent nuclei from the lateral surface. We quantified 54.41 ± 2.99 % (**, p<0.01) of nuclei adjacent to the lateral surface and 23.21 ± 2.94 % (***, p<0.001) of non-adjacent nuclei. The average distance of ventricular nuclei was also increased by 20.52 ± 2.59 % (***, p<0.001). Injection of Nrp2-Fc and Sema3B-Fc had no impact on the radial positioning of the PH3-reactive cells. We detected 65.33 ± 2.12 % (p=0.59) nuclei directly next to the lateral surface, 10.64 ± 1.66 % (p=0.6) of non-adjacent nuclei. However, the distance of ventricular nuclei was significantly decreased by 18.91 ± 2.43 % (***, p<0.001) in comparison the control-Fc. These experiments show that manipulation of the Sema/Nrp-signalling could alter the radial dispersion of mitotic nuclei positioning in the ventricular zone in a short range of time, consistent with an implication of endogenous CSF-derived Sema3s/Nrps complexes in regulating apical nuclei position *in vivo*. To further validate these findings, we designed an *ex vivo* experimental paradigm to visualize Sema/Nrp-induced rapid changing of apical nuclei positioning. Cortical tissue was dissected and incubated for 1 hiv with the apical surface apposed to coverslips coated with Sema3F-Fc/Nrp1-Fc, Sema3B-Fc/Nrp2-Fc, or control-Fc, respectively. Staining of F-Actin allowed us to visualize the apical endfeet and to control the integrity of the tissue. We quantified the apical density of nuclei (Figure 5D). In the Sema3F-Fc/Nrp1-Fc condition, the proportion of nuclei on the apical surface was decreased by 23.21 ± 2.94 % (***, p<0.001) in comparison to control-Fc. In contrast, in the Sema3B-Fc/Nrp2-Fc condition, it was significantly increased by 13.85 ± 4.12 % (*, p<0.05) compared to control-Fc. Thus, Sema3B/Nrp2 and Sema3F/Nrp1-signalling exert apical and anti-apical forces, which regulate the positioning of mitotic nuclei along the cortical baso-apical axis.

We further investigated whether these nuclear pushing and pulling forces influence the timing during which apical progenitor cells reside at the ventricular surface during division. To address this question, we developed a sophisticated *en face* imaging of cortical cultures, where we monitored the apical nuclei of cortical tissue *ex vivo*. Cortices of E13.5 embryos were dissected and the ventricular side apposed to glass bottom dishes coated with Sema3F-Fc/Nrp1-Fc, Sema3B-Fc/Nrp2-Fc or control-Fc, respectively. The brains were incubated with Syto16 for nuclear staining and SirActin to visualize the apical endfeet and monitoring the integrity of the tissue (Figure 5E). In line with the previous experiments, we observed differences in nuclear density on the apical side. In the Sema3F-Fc/Nrp1-Fc condition, numbers of apical nuclei were significantly decreased by 10.97 ± 2.54 % (***, p<0.001) compared to control-Fc. In contrast, tissue apposed to Sema3B-Fc/Nrp2-Fc revealed an increase of apical nuclei by 12.29 ± 5.97 % (p=0.058), once more suggesting a role of Sema/Nrp-interactions on apical mitotic nuclei positioning.

We next analysed the duration of individual nuclei stay at the apical surface and the timing of the different cell cycle phases (Figure 5F). Incubating the cortices on control-Fc resulted in an overall time of stay of about 188.22 ± 8.39 min. This duration was subdivided in 68 ± 5.6 min for the appearance and prophase, 53 ± 3.4 min for the metaphase, and 67 ± 3.4 min for ana-/telophase and disappearance from the ventricular surface. Strikingly, in the presence of Sema3F-Fc/Nrp1-Fc, apical nuclei spent just 91.85 ± 3.97 min on the apical surface, thus almost half of the time of controls (26 ± 1.6 min appearance and prophase, 34 ± 2.1 min metaphase, and 32 ± 1.4 min for ana-/telophase and disappearance). In the Sema3B-Fc/Nrp2-Fc condition, apical nuclei stayed 147.1 ± 7.11 min (***, p<0.001) at the ventricular surface (38 ± 3.3 min appearance and prophase, 54 ± 3.9 min metaphase, and 55 ± 3.6 min for ana-/telophase and disappearance). Notably, the exposure to extrinsic Sema/Nrp-molecules resulted in an accelerated mitosis at the apical surface, although Sema3B/Nrp2- and Sema3F/Nrp1-signalling had different outcomes. These results show that the pro- and anti-apical forces exerted by apical Sema/Nrp-signalling influence the time of apical stay of cortical progenitors and the duration of the cell cycle phases.

### The generation of intermediate precursors and postmitotic neurons are modulated by CSF-derived Semas/Nrps

The regulation of apical adhesion and nuclear positioning is particularly important during asymmetric cell division, when one daughter cell remains attached in the neuroepithelium as an apical progenitor, whereas the other one disengages from the ventricular surface to differentiate into a neuron or an intermediate progenitor, which then gives rise to two neurons. The inactivation of GSK3 in cortical progenitor cells was reported to result in enhanced cortical proliferation whereas GSK3-activation promotes neuronal differentiation ^39,40^. This broaches the issue whether CSF-derived Sema and Nrps are implicated in the control of fate determination of apical progenitor daughter cells in the VZ. To address this question, we analysed whether deletion of Semas and Nrps affects the number and distribution of intermediate precursor cells. Embryonic coronal brain slices of E12.5 old wildtype and mutant embryos were stained with an antibody against Tbr2 (T-brain 2), a specific molecular marker to identify transient amplifying cells ^41–43^, and Tbr2-positive cells were counted (Figure 6A,C). We found decreased numbers of IPCs in Sema3F -/- and Nrp1^Sema/Sema^ mice in comparison to wildtype littermates of about 18.02 ± 5.97 % (*, p=0.037) and 13.64 ± 3.39 % (**, p=0.003), respectively. In contrast, Sema3B and Nrp2 knockout embryos showed significant augmented numbers of transient amplifying cells. Sema3B -/- mice reveal 76 ± 4.25 % (***, p<0.001) more IPCs and in Nrp2 deficient mice the numbers were increased for about 32.65 ± 4.12 % (***, p<0.001). Transient amplifying cells are the most prominent precursors during middle and late neurogenesis ^44^ and they generate the majority of postmitotic neurons of the cerebral cortex ^45,46^. However, during early neurogenesis, the majority of postmitotic neurons are produced directly by apical progenitor cells ^46^. Therefore, we further investigated the numbers of postmitotic neurons generated in the different mouse lines at E12.5, using an antibody against Tbr1 (T-brain 1) that is specifically expressed in early-born neurons and of layer 6 ^41,43^. In agreement with our previous findings of the intermediate progenitor pool, Sema3F -/- and Nrp1^Sema/Sema^ embryos exhibited lower numbers of postmitotic neurons at E12.5 (Figure 6B,D). In comparison to wildtype littermates, decrease by 20.34 ± 5.28 % was observed in Sema3F -/- embryos (**, p=0.009) and by 34.51 ± 7.39 % (***, p<0.001) in Nrp1^Sema/Sema^ embryos. In contrast, the number of postmitotic neurons was increased by 46.48 ± 3.64 % (***, p<0.001) and 22.36 ± 3.2 % (***, p<0.001) in Sema3B -/- and Nrp2 -/- embryos, respectively. These results indicate that the pro-apical Sema3B/Nrp2-signaling inhibits the generation of Tbr1-positive neurons and Tbr2-positive intermediate progenitor cells, whereas the anti-apical Sema3F/Nrp1-signalling promotes differentiation.

**Figure 6:**
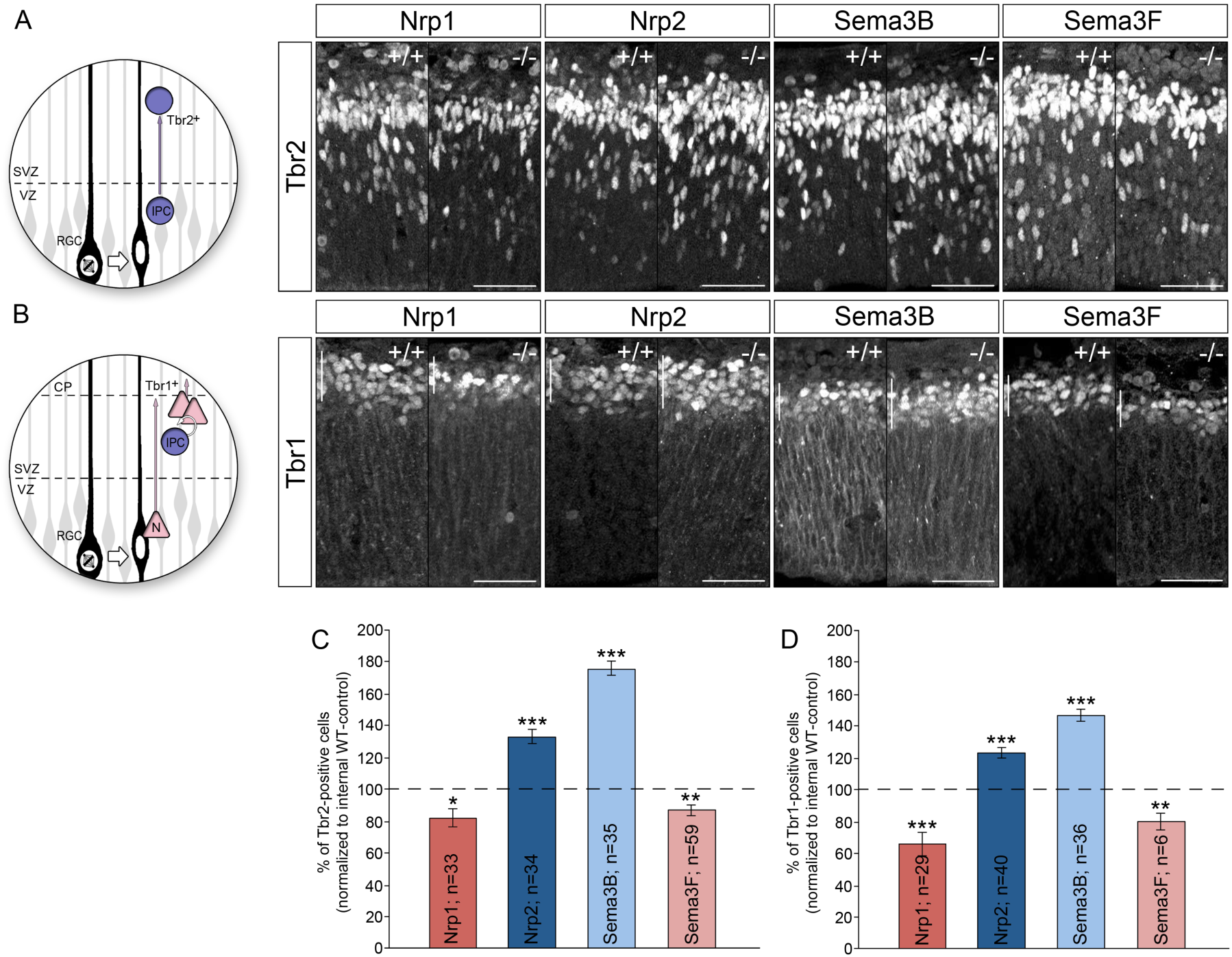
Sema/Nrp-complexes regulate the generation of intermediate precursors and postmitotic cortical neurons. **(A)** Tbr2-positive intermediate progenitors delaminate from the apical surface and translocate their soma into the SVZ. Microphotographs illustrate Tbr2-positive nuclei in wildtype, Nrp1^Sema/Sema^, Nrp2 -/-, Sema3B -/-, and Sema3F -/- embryonic cortices. **(B)** During early neurogenesis, Tbr1-positive postmitotic neurons are generated directly by radial glia cells or indirectly by intermediate progenitor cells. Microphotographs illustrate Tbr1-positive nuclei in Nrp1^Sema/Sema^, Nrp2 -/-, Sema3B-/-, and Sema3F -/-embryonic cortices. **(C)** The histogram shows that Sema3F -/- and Nrp1^Sema/Sema^ cortices have less Tbr2-positive cells in comparison to the wildtype littermates at E12.5. In contrast, Sema3B and Nrp2 knockout embryos show significant augmented numbers of intermediate progenitor cells. **(D)** The histogram shows that Sema3F -/- and Nrp1^Sema/Sema^ mice exhibit reduced numbers of Tbr1-positive neurons in the cortical plate at E12.5. In contrast, Tbr1-reactive cells are increased in Sema3B and Nrp2 knockouts, in comparison to WT littermates. Scale bars: 50 µm; paired t-test, * p<0.05, ** p<0.01, *** p<0.001.

## Discussion

Our study provides evidence for an unconventional Sema/Nrp-signalling mechanism in which both, ligands and receptors are synthesized by the nascent choroid plexus and released into the CSF. Sema3s represent a class of secreted molecules and for Sema3B and Sema3F an expression in the developing choroid plexus have already been reported ^15,47^. To our surprise, we found that Nrp1 and Nrp2, which are, apart for Sema3E, obligatory components of Sema3s receptors, are not expressed by apical progenitors. Rather, they are produced by the nascent choroid plexus, the floor plate and meninges, tissues known to secrete molecules in the CSF. Soluble forms of Nrps have been previously reported in physiological as well as pathological contexts ^20^. Particularly, Nrp1 in the CSF was proposed to be associated with Alzheimer disease and aging in humans ^48,49^. Soluble forms of Nrps in the CSF could arise from splice variants encoding forms devoid of transmembrane domains, like those previously described for both Nrp1 and Nrp2 ^19,50^. Soluble forms of Nrps can also result from ectodomain shedding ^51^. Up to now, the biological functions of these soluble Nrps has remained elusive. *In vitro* application of soluble Nrps have various outcomes which might depend on the expression profile of Nrp binding partners and components of downstream signalling cascades in the exposed cells. Hence, soluble Nrps were found to display dominant negative effects on the Sema3 signalling, acting via ligand titration and competition with endogenous Nrps. The Nrp1-extracellular domain was also found to reverse Sema3E/PlxnD1-mediated repulsion to attraction in subiculo-mamillary neurons ^52^. Our findings provide the first evidence that *trans* delivery of soluble Nrps can reconstitute Sema3 receptor complexes with dual functions in neural progenitors expressing Plexins only. The conformation of Sema/Plxn/Nrp-complexes has been recently highlighted by crystallography data reporting that Nrp ectodomain is indeed required for stabilizing weak interactions contracted between Sema3 and PlxnA molecules ^53^. Interestingly, these structural features are fully compatible with the mode of presentation of the different components of the ternary complex that we report in the developing cerebral cortex, with the CSF delivering pre-formed Sema3/Nrp-complexes to apical Plxns. We found that apical progenitor cells express Plexins, particularly PlxnB1, that accumulate at the ventricular surface of progenitor cells, the expected location for interactions with CSF-derived Sema/Nrp complexes. Contributions of Plxnb1/2 in the CSF-derived Sema3B and Sema3F signalling is strongly supported by previous work reporting that Plxnb1/b2 double mutant mouse embryos exhibit cortical development defects, with decreased proliferation and reduced neuronal production, resulting in cortical thinning ^23^. PlxnB1 and PlxnB2 are likely redundant since individual deletions do not strongly affect cortical development ^23,54–56^. Interactions between Sema3B/3F, Nrp1/2 and PlxnAs have been demonstrated in several contexts ^57^. In contrast, Sema3B/3F-binding to PlxnB co-receptors via transmembrane Nrps has not been reported so far, and could be specific to this particular *cis* and *trans* configuration of ligand/receptor-components.

We found that two conspicuous combinations of Sema3/Nrps, Sema3B/Nrp2-complexes and Sema3F/Nrp1-complexes exert opposite influences on cortical progenitor adhesion and positioning. Whereas opposite outcomes of Sema/Nrp/Plxn-signalling have been observed in various contexts ^58^, the generation of duality of the effects remains puzzling. First, differences might come from specific conformations of ternary Sema3/Nrp/Plxn-complexes, possibly raised from binding interactions, but also from secondary interfaces formed between three-dimensional domains, reported to regulate the affinity and specificity of modular domain-mediated interactions ^59^. Second, differences could already be prefigured by Sema3/Nrp-complexes, since prominent preferential association of Sema3B with Nrp2 rather than Nrp1 was observed in our biochemical analysis of complexes in the CSF of Sema3B-GFP ki embryos. Finally, Sema3B/Nrp2-complexes and Sema3F/Nrp1-complexes could preferentially associate with distinct Plxn co-receptors, then triggering specific downstream signalling and functional outcomes. Deep investigations of the specificities generated by these different architectures of ligand/receptor-complexes are indispensable for further understanding the variety of biological outcomes they generate.

Our data strongly suggest that this dual Sema/Nrp-signalling regulates the dynamic of apical progenitor cells. With our loss-of-function and gain-of-function experiments, we show that the Sema3B/Nrp2-couple favours the apical accumulation of nuclei in prophase and anaphase, whereas the Sema3F/Nrp2-complex allows a sub-apical positioning of nuclei. Second, our timelapse analyses show that the mitosis duration is modified upon Sema3s exposure. These results thus reveal that the Sema3/Nrp-signalling regulates both, the distance of nuclei to the ventricular surface and the apical residence time. CSF-derived cues were shown to regulate progenitor proliferation and differentiation ^2,60^. Interestingly, we found that the generation of neurons and intermediate progenitors is increased in Sema3B and Nrp2 mutants, whereas it is decreased in Sema3F and Nrp1 mutants. We thus propose that the influences of CSF-derived dual Sema/Nrp-signalling on apical progenitor dynamics are important for cell fate determination of the progeny. This view is supported by previous studies indicating that neurogenesis progress is tightly linked to the interkinetic nuclear migration (INM). Apical progenitor cells undergo interkinetic nuclear migration along the apico-basal axis and disruption of the apico-basal nuclei movement results in premature depletion of cortical progenitor cells and increased neurogenesis ^61^. In support, the amplitude of the INM was proposed to be predictive for the neurogenic nature of the next division ^62^. In addition, apico-basal gradients of neurogenic and proliferative signalling have been reported to influence neural progenitors according to the position of their nuclei during INM ^63–66^. Likewise, nuclei located at different apico-basal positions would be exposed to different concentrations of these signals and therefore might acquire distinct proliferative and neurogenic features ^67^. Apical position also correlates with cell fate in other biological systems. For example, in the zebrafish spinal cord live cell imaging of asymmetric progenitor divisions revealed that the more apical daughter cell inheriting the apical Par3-domain becomes a neuron, whereas the cell inheriting either the basal process or molecular components associated with the basal process retains progenitor fate ^68^.

Our timelapse imaging revealed that exposing apical progenitor cells to distinct Sema/Nrp-combinations result in changes in the residence time at the apical surface. We observed that both combinations of Sema/Nrp-complexes significantly shortened the overall progenitor apical stay. Although having opposite effects on adhesion, the two complexes did not alter in opposite way the duration of apical stay suggesting that any type of imbalanced apical attachment is reflected in mitosis shortening. Because removal of the CSF resulting from cortex opening prevents apical progenitor access to many signals, it is possible that some of them lacking in our set-up act complementary to Sema/Nrps to regulate the cell cycle length. Notably, Sema3F/Nrp1-signalling had a striking pronounced effect, in particular reflected in a marked reduction of the metaphase, while this effect was not observed in the Sema3B/Nrp2 condition. Interesting links are suggested between the duration of apical stay and cell fate decisions. In brain organoids, the length of the prophase/metaphase was found to distinguish human and non-human primate apical progenitors. Moreover, the metaphase was found to be shorter for cells committed to become neurons than for those keeping a progenitor fate ^69^. In addition, previous studies showed that treatment leading to a mitotic delay also alters the fate of RGC progeny ^70^. These data suggest that the modulation of mitotic progress by Sema/Nrp-complexes could be a mechanism by which these signals influence cell fate decisions during corticogenesis.

Our study also shows that dual Sema3/Nrp-signals oppositely control the ability of cortical progenitors to settle on a substrate *in vitro*. Interestingly, manipulations of the Eph/Ephrin-signalling in neural progenitor cells were found to alter apical adhesions, resulting in displaced nuclei and changes of cell fate during the neurogenic period ^37^. Our findings raise the question of how these dual Sema3-mediated properties contribute to nuclear positioning in the neuroepithelium. Adhesiveness of cells on a substrate *in vitro* depends on actin cytoskeleton assembly and coupling with transmembrane adhesion proteins, as well as on internal relaxing versus contractile forces generated by molecular motors. In the context of cell guidance, attractive or repulsive activities of Semaphorins have been shown to rely on these mechanisms ^71,72^. Interestingly, internal contractility within the apical domain is also thought to be an important parameter of progenitor divisions. The apical domain, which is composed of adherent junctions and a cortical actin network, is remodelled during the INM, enlarging when nuclei are close to the ventricular surface and shrinking when they migrate basally ^73^. In addition, Myosin activity, which impacts apical actin organisation, is crucial for nuclear positioning ^61^. Thus, a Sema3/Nrp-mediated program could orchestrate the morphological remodelling of progenitor cells and relocation of their nuclei during neurogenesis progression by regulating internal contractile properties at their apical pole.

Moreover, modulation of the adhesion of apical feet is crucial to the delamination of neural progenitors from the ventricular surface and is thought to be an instructive event of the emergence of basal progenitors and neurons ^7,9^. Apically-delivered Sema3/Nrp-cues could thus modulate the generation of intermediate progenitors and neurons through the control of neural progenitor adhesion of apical endfeet, which would be consistent with reported functions of Semaphorins. Beyond their contribution to the pathfinding of axonal growth cones during neural circuit formation, axon guidance molecules also play prominent roles in cell shape and adhesion remodelling accompanying tissue formation and function. For example, the Semaphorin-signalling regulates morphological remodelling of podocytes in the kidney ^74^ and hypothalamic neurons secreting gonadotropin-releasing hormone (GnRH) in the brain ^75^.

Interestingly, the release of various additional axon guidance molecules in the CSF, such as Sema4D, Sema7A, Sema3C, and Slits, was already reported ^16,76,77^. Our findings bring the first insights into a likely much broader developmental program orchestrating apical dynamics of cortical progenitors. Moreover, wether the contributions of such apically delivered axon guidance molecules have been extended through evolution is a fascinating question for future investigations, given the great diversity of cortical progenitor shapes and radial anchors that emerged in the primate lineage ^1,78,79^.

## Acknowledgments

We thank Christiana Rurhberg and Laura Denti for providing us with Sema3F embryos. We thank Julien Falk for technical advices, Christiana Rurhberg for scientific discussions, and Edmund Derrington for critical reading of the manuscript. This work was supported by ERC (European Research Council) grant 281604-YODA to VC. It was conducted within the frame of the LabEx CORTEX and DEVWECAN of Université de Lyon, within the program ‘‘Investissements d’Avenir’’ (ANR-11-IDEX-0007) operated by the French National Research Agency (ANR).

## METHODS

### Animals

All animal procedures were performed in accordance to European Communities Council Directive and approved by French ethical committees. Time pregnant mice of following strains were used: Sema3Bko ^14^, Sema3Fko ^24^, Nrp2ko ^25^, and Nrp1^Sema/Sema^ mice in which the Sema3 binding is selectively disrupted ^26^. The day of insemination was considered as embryonic day 0.5 (E0.5). Mice were bred and maintained under standard conditions with access to food and water ad libitum on a 12 h light/dark cycle.

### Section preparation

For the preparation of time-staged embryonic brains time pregnant mice were deeply anesthetised with 10 % chloral hydrate and the embryos dissected. Embryonic heads were fixed over-night in 4 % PFA (in 1X PBS, pH 7.4) at 4 °C, followed by a sequential Sucrose treatment (10 %, 15 %, 30 % in 1X PBS, pH 7.4) as cryoprotection. After freezing in Isopentan and dry ice at – 40 °C, heads were sectioned into 20 µm slices (at -21 °C) using a Cryotome CM1900 (Leica) and mounted on Superfrost Plus slides (Thermo Fisher Scientific).

### Immunolabelling

Brain slices, cortical preparations, and dissociated single cells were washed in 1X PBS (pH 7.4) with 0.2 % Tween20, followed by blocking (in 4 % BSA in 1X PBS/0.2 % Tween20) for 2h at RT and an incubation with the primary antibody over night at 4 °C. After washing, the secondary antibody was applied for 2 h at RT. The nuclei were stained for 15 min at RT with Hoechst33342 (1 ng/ml in H_2_O). As primary antibodies were used: rabbit anti-GFP (Invitrogen, 1:100), goat anti-Nrp1 (R&D, 1:100), goat anti-Nrp2 (R&D, 1:100), rat anti-PH3 (Sigma, 1:50), mouse anti-γTubulin (Sigma, 1:100), rabbit anti-PlexinA2 (Santa-Cruz, 1:25), rabbit anti-PlexinB1 (Abcam, 1:100), rat anti-PlexinB2 (Invitrogen, 1:100), rabbit anti-Alcaline Phosphatase (Gene hunter, 1:300), rabbit anti-phospho-GSK-3α/β (Ser21/9) (Abcam, 1:100), rabbit anti-Sox2 (Abcam, 1:400), rabbit anti-Tbr1 (Abcam, 1:400), rabbit anti-Tbr2 (Abcam, 1:400). As secondary antibodies were used: donkey anti-goat IgG Alexa488 (Molecular Probes, 1:500), donkey anti-goat IgG Alexa555 (Molecular Probes, 1:500), donkey anti-mouse IgG FP547H (Interchim, 1:500), donkey anti-rabbit IgG FP647H (Interchim, 1:500), donkey anti-rabbit IgG Alexa488 (Inv, 1:500), donkey anti-rat IgG Alexa488 (Molecular Probes, 1:500), donkey anti-rat IgG FP647H (Interchim, 1:500).

### *In-situ* Hybridisation

Digoxigenin (Dig)-labelled probes against Semaphorin3B, Semaphorin 3F Nrp1, and Nrp2 were used as previously described ^68^. In brief, sectioned were fixed for 10 min in 4 % PFA (in 1X PBS, pH 7.4), permeabilised for 10 min in 0.2 M HCl and acetylated in 0.1 M triethanolamine with 5 mM acetan hydrid for 15 min. Hybridisation were performed over night at 56 °C with a probe concentration of 3 ng/μl. Slides were blocked for 2–3 h, using 2 % blocking reagent (Roche), followed by the detection of the Dig-labelled riboprobe with an anti-DIG Fab fragment conjugated with alkaline phosphatase (1:750; Roche). The colometric reaction was performed using mixture of 5-Bromo-4-chloro-3-indolyl phosphate (Roche) and Nitro blue tetrazolium chloride (Roche).

### Western blotting and immunoprecipitation

CSF was directly denatured for 5min at 95 °C in Laemmli buffer. Cortical tissue was lysed for 45 min on ice in 0.5 % NP-40 (Sigma-Aldrich), 0.5 % SDS, 150 mM NaCl, 50 mM Tris-HCl, 2 mM EDTA, and protease inhibitor cocktail (Roche). For western blotting samples were denaturated for 5 min at 95 °C in Laemmli buffer. For immunoprecipitation, lysates were incubated at 4 °C overnight with rabbit anti-GFP (1:1000, Roche) and pull down was performed with magnetic proteinA beads (Millipore) for 15 min at RT. After several washing steps, proteins were collected in 80 ml Laemmli buffer and 20 ml of solution was used for SDS-Page (10 % acrylamide precast mini gels, Bio-Rad). A Trans-Blot Turbo Transfer System (Bio-Rad) was used to transfer the proteins in a nitrocellulose membrane that was blocked in 5 % milk powder (in 1X PBS/0.2 % Tween20) for 30 min at RT and incubated with the primary antibody over night at 4 °C. Membranes were washed in 1X PBS/0.2 % Tween20 and incubated with the secondary antibody for 1 h at RT. For signal detection, ECL Prime was used according to the manufacture instructions (RPN2232, G.E. healthcare) and the chemidoc MP system with applied software (Bio-Rad). As primary antibodies were used: rabbit anti-GFP (Invitrogen, 1:5000), rabbit anti-phospho-GSK-3α/β (Ser21/9) (Abcam, 1:500) goat anti-Nrp1 (R&D, 1:1000), and goat anti-Nrp2 (R&D, 1:1000). As secondary antibodies were used: anti-goat HRP (Sigma-Aldrich, 1:5000) and anti-rabbit HRP (Sigma-Aldrich, 1:5000).

### Preparation of cortical single cells

Time pregnant mice were deeply anesthetised with 10 % chloral hydrate. Brains of the embryos were dissected in PBS (Invitrogen) supplemented with 0.65 % glucose, cortical tissue was dissected and incubated in PBS with 2.5 % trypsin for 17 min at 37 °C. Afterwards, tissue was dissociated by titruation and filtered through nylon gaze to remove cell aggregates. Cells were seeded on coverslips coated with 19.5 µg/ml Laminin (Sigma-Aldrich) and 5 µg/ml Poly-L-Lysin (PLL, Invitrogen) at a density of 300 cells/mm^2^ and incubated at 37 °C and 5 % CO2 in a humid atmosphere in DMEM (Invitrogen) supplemented with 10% FBS, 10000 U/ml penicillin, 10000 μg/ml streptomycin, 0.065 % D-glucose and 0.4 mM L-glutamine. For immunostaining, cells were fixed in 4 % PFA (in 1X PBS, pH 7.4) for 10 min at room temperature.

### Binding assay

To test the binding of recombinant Semaphorin/Neuropilin complexes to apical progenitors, we incubated brain hemispheres with recombinant Sema3B-Alkaline Phosphatase in the presence or absence of recombinant Nrp1-Fc or Nrp2-Fc (R&D systems), respectively (10 ng/µl in PBS). After 30 min of incubation at 37 °C and 5 % CO2 in a humid atmosphere in DMEM (Invitrogen), hemispheres were fixed in 4% PFA (in 1X PBS, pH 7.4) over night at room temperature and immunolabelling was performed as described, followed by dissection of the somatosensory cortex and flat mounting with Mowiol.

### *En face* life cell imaging

Time pregnant mice were deeply anesthetised with 10 % chloral hydrate. Brains of the embryos were dissected in ice cold PBS (Invitrogen) supplemented with 0.65 % glucose. The hemispheres were separated and incubated with Syto16 (Invitrogen, 1:1000) and SirActin (Spirochrome, 1:1000) in DMEM (Invitrogen) for 20 min on ice. After transferring the hemispheres to ice cold DMEM, the somatosensory cortex was dissected and placed into glass bottom dishes (Matec) that were coated with 1 µg/ml Poly-L-Lysin and recombinant proteins in different combinations (10 ng each) for 1 h at 37 °C. After incubation for 20 min at RT in 100 µl culture medium (DMEM (Invitrogen) with 25 % HBSS (Invitrogen), 10% FBS, 10000 U/ml penicillin, 10000 μg/ml streptomycin, 0.65 % D-glucose and 0.4 mM L-glutamine) supplemented with 0.5 % methyle cellulose, compartments were filled with culture medium and 0.1 mM HEPES (Invitrogen) before live cell imaging was performed. Pictures were taken at 10 min interval for 10 h, using IQ3 software with multi-positions and Z-stack protocols. To reduce exposure time and laser intensity acquisition were done using binning 1×1.

### Detection and analysis

Pictures of *in situ* hybridization experiments were taken using a Z1 observer microscope (Zeiss). Pictures of the apical surface or brain slices were taken with an inverted confocal laser-scanning microscope FV1000 (Olympus). *En face* live cell imaging was performed using the Olympus IX81 microscope equipped with a spinning disk (CSU-X1 5000 rpm, Yokogawa), an Okolab environmental chamber, an EMCCD camera (iXon3 DU-885), and applied software (Andor technology). Photographs were analyzed with Fiji software ^69^. Data collection and analysis was performed blind except for the *en-face* imaging. Analysis of cell division orientation was performed as described in Arbeille et al. (2015). Results represent at least three independent experiments or three analysed brains out of two litters (except in-vivo injection in the lateral ventricle, were results represent two independent experiments). Quantifications are represented as mean ± SEM and statistical significance was tested as indicated in the Figure legends. * p<0.05 was considered as significant, ** p<0.01, and *** p<0.001.

## Figure legends

**Figure S1:**
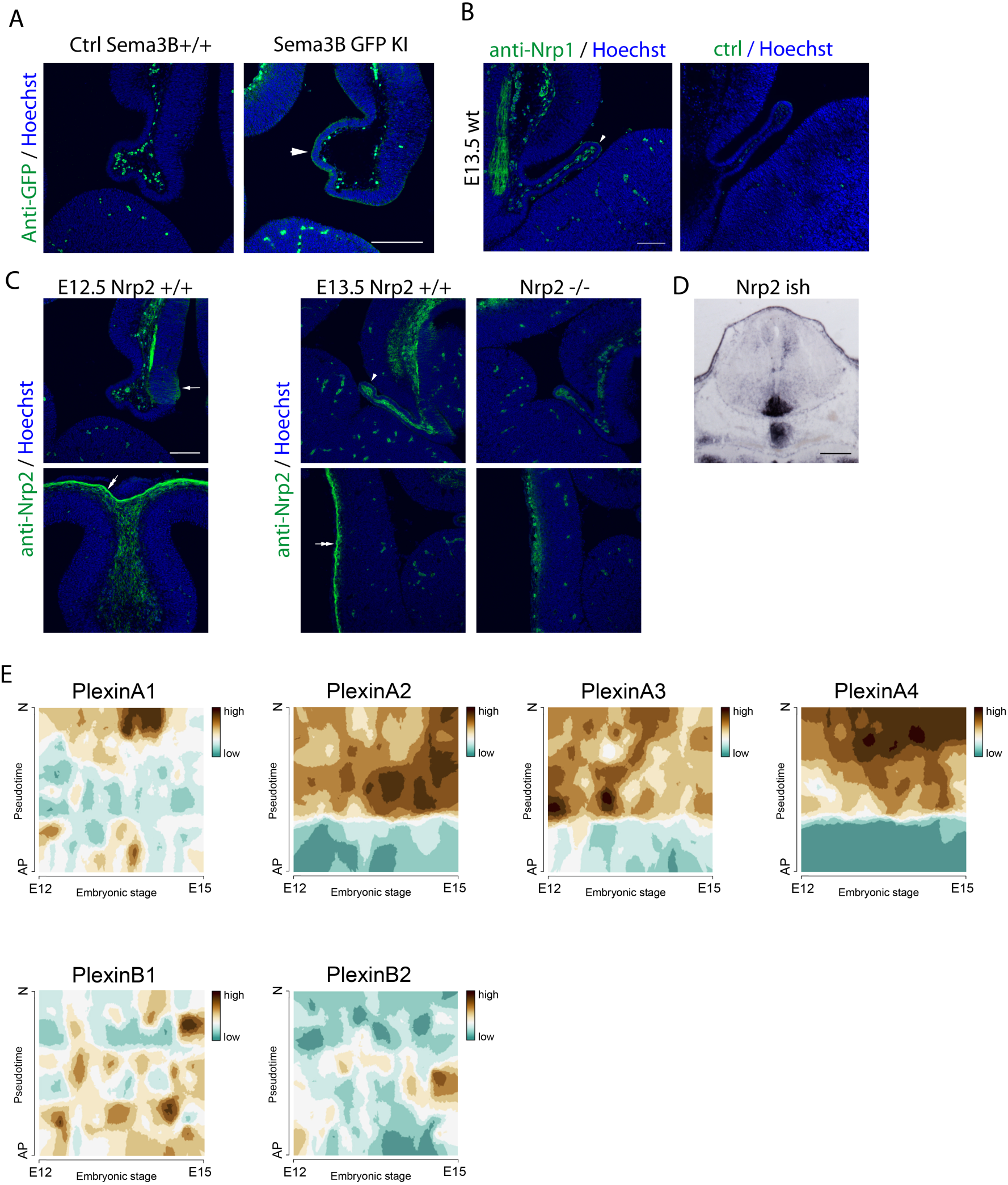
Sources of Semas, Nrps and Plexins in the embryonic brain. **(A)** GFP-immunolabeling of Sema3B-GFP ki/ki and control embryonic brains show the presence of Sema3B proteins at the apical border of the telencephalic choroid plexus at E12.5. **(B)** Microphotographs illustrating the control of Nrp1 labelling at the choroid plexus apical border by comparative immunostaining using secondary antibody alone. **(C)** Anti-Nrp2 immunolabeling of E12.5 and E13.5 embryonic brains and labelling of E13.5 Nrp2 ^-/-^ brain sections show Nrp2-labelling specificity. Arrowhead point to the telencephalic choroid plexus, arrow point to the diencephalic ventricular zone and double arrow point to the meninges. Note the enrichment of Sema3B-GFP, Nrp1 and Nrp2 at the apical border of CP epithelial cells. **(D)** *In situ* hybridization detects abundant Nrp2 transcripts in the midbrain floor plate at E13.5. **(E)** Single cell transcriptome analysis of embryonic cortical cells reveals the expression of Plexins at different embryonic stages (between E12-E15). Pseudotime describes the differentiation from an apical progenitor cell (0) to a postmitotic neuron (1). Scale bar=100µm

**Figure S2:**
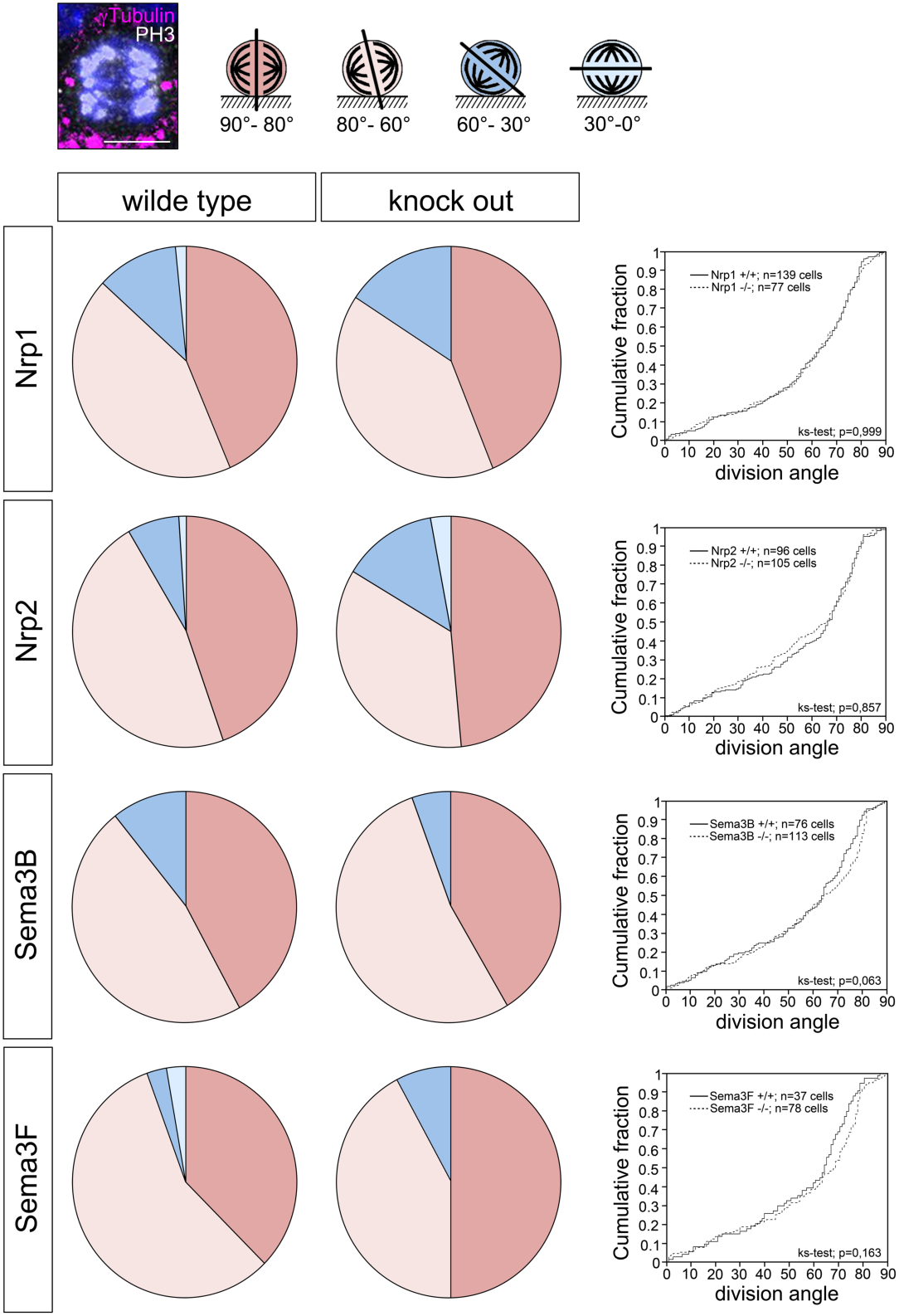
CSF-derived Sema/Nrp-complexes are dispensable for progenitor division orientation. E12.5 cortical sections of WT, Sema3F -/-, Nrp1^Sema/Sema^, Sema3B -/- and Nrp2 -/- embryos were stained with antibodies against phospho-histone 3 (PH3) and γTubulin and the division orientation of cells in ana-/telophase with respect to the apico-basal axis was examined according to Arbeille et al. (2015). Angle distributions are represented in distinct classes (90°-80°, 80°-60°, 60°-30°,30°-0) in pie charts and as cumulative fraction curves. No significant differences in the orientation of the division plane was observed. Scale bar 5 µm; n= number of analysed cells.

**Figure S3:**
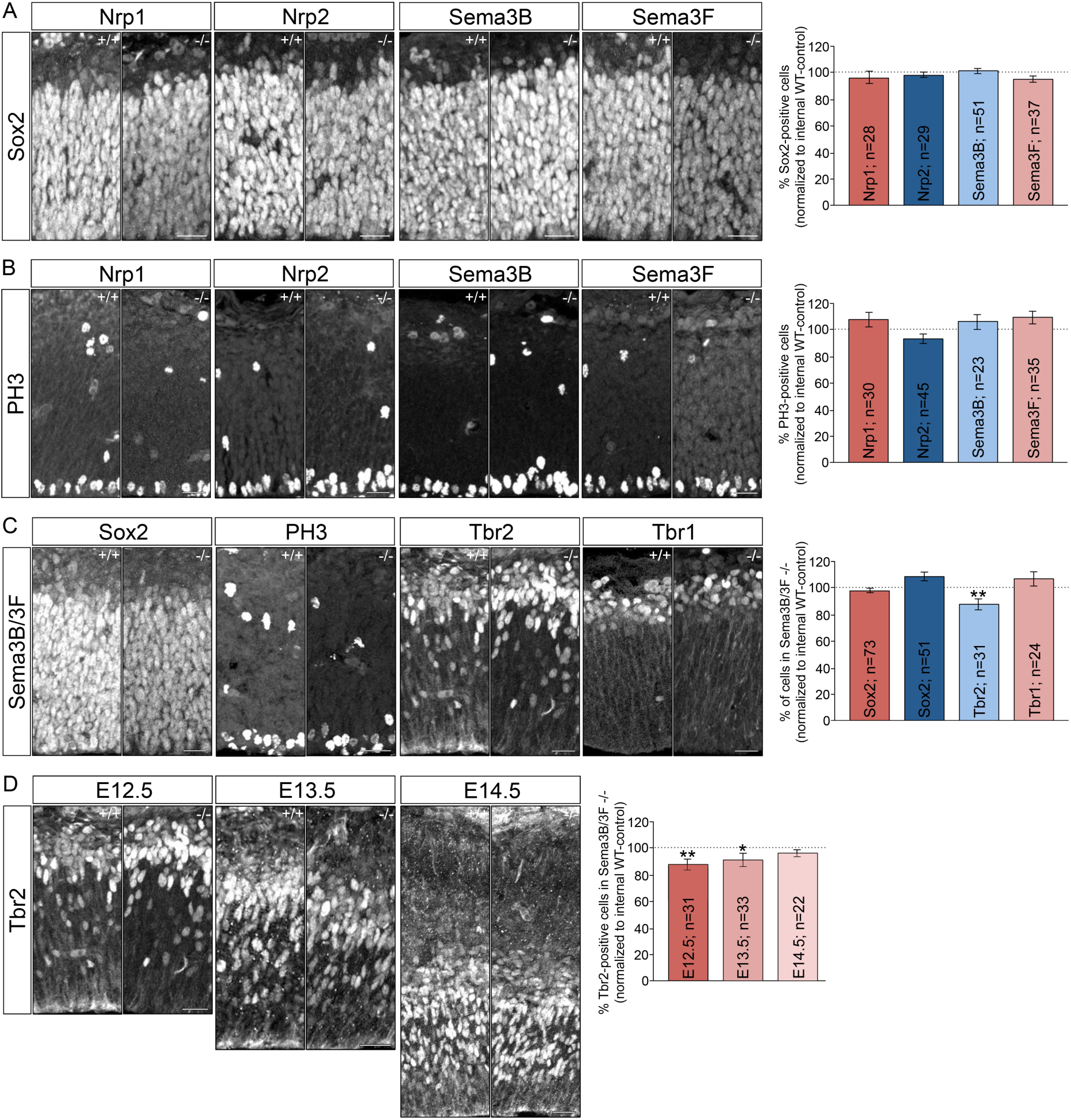
The number of Sox2 and PH3-positive cells are not altered in the absence of CSF-derived Sema/Nrp-molecules. E12.5 cortical sections of WT, Sema3F -/-, Nrp1^Sema/Sema^, Sema3B -/- and Nrp2 -/- embryos were stained with antibodies against **(A)** Sox2 (sex-determining region Y-box containing gen 2) and **(B)** phospho-histone 3 (PH3) and positive nuclei were counted. No significant changes were observed. **(C)** Immunolabelling of Sema3B/3F double mutant embryos against Sox2, PH3, and Tbr1 do not reveal significant changes in comparison to wildtype littermates. However, the numbers of Tbr2-positive cells are reduced in Sema3B/3F knockout mice. This effect disappears at later stages (**D**). Scale bars 25 µm; n= number of analysed sections.

## References

1. Gotz, M. & Huttner, W. B. The cell biology of neurogenesis. Nat. Rev. Mol. Cell Biol. 6, 777–788 (2005).

2. Lehtinen, M. K. & Walsh, C. A. Neurogenesis at the brain-cerebrospinal fluid interface. Annu. Rev. Cell Dev. Biol. 27, 653–679 (2011).

3. McConnell, S. K. Constructing the cerebral cortex: neurogenesis and fate determination. Neuron 15, 761–768 (1995).

4. Rakic, P. A small step for the cell, a giant leap for mankind: a hypothesis of neocortical expansion during evolution. Trends Neurosci. 18, 383–388 (1995).

5. Gloerich, M., Bianchini, J. M., Siemers, K. A., Cohen, D. J. & Nelson, W. J. Cell division orientation is coupled to cell-cell adhesion by the E-cadherin/LGN complex. Nat. Commun. 8, 13996 (2017).

6. Miyamoto, Y., Sakane, F. & Hashimoto, K. N-cadherin-based adherens junction regulates the maintenance, proliferation, and differentiation of neural progenitor cells during development. Cell Adh. Migr. 9, 183–192 (2015).

7. Zhang, J. et al. Cortical neural precursors inhibit their own differentiation via N-cadherin maintenance of beta-catenin signaling. Dev. Cell 18, 472–479 (2010).

8. Wang, Q., Yang, L., Alexander, C. & Temple, S. The niche factor syndecan-1 regulates the maintenance and proliferation of neural progenitor cells during mammalian cortical development. PLoS One 7, e42883 (2012).

9. Das, R. M. & Storey, K. G. Apical abscission alters cell polarity and dismantles the primary cilium during neurogenesis. Science 343, 200–204 (2014).

10. Paridaen, J. T. M. L. & Huttner, W. B. Neurogenesis during development of the vertebrate central nervous system. EMBO Rep. 15, 351–364 (2014).

11. Gerstmann, K. et al. Thalamic afferents influence cortical progenitors via ephrin A5-EphA4 interactions. Development 142, 140–150 (2015).

12. Borrell, V. et al. Slit/Robo signaling modulates the proliferation of central nervous system progenitors. Neuron 76, 338–352 (2012).

13. Qiu, R. et al. Regulation of neural progenitor cell state by ephrin-B. J. Cell Biol. 181, 973–983 (2008).

14. Falk, J. et al. Dual functional activity of semaphorin 3B is required for positioning the anterior commissure. Neuron 48, 63–75 (2005).

15. Arbeille, E. et al. Cerebrospinal fluid-derived Semaphorin3B orients neuroepithelial cell divisions in the apicobasal axis. Nat. Commun. 6, 6366 (2015).

16. Johansson, P. A. The choroid plexuses and their impact on developmental neurogenesis. Front. Neurosci. 8, 340 (2014).

17. Telley, L. et al. Sequential transcriptional waves direct the differentiation of newborn neurons in the mouse neocortex. Science 351, 1443–1446 (2016).

18. Raimondi, C. & Ruhrberg, C. Neuropilin signalling in vessels, neurons and tumours. Semin. Cell Dev. Biol. 24, 172–178 (2013).

19. Rossignol, M., Gagnon, M. L. & Klagsbrun, M. Genomic organization of human neuropilin-1 and neuropilin-2 genes: identification and distribution of splice variants and soluble isoforms. Genomics 70, 211–222 (2000).

20. Gagnon, M. L. et al. Identification of a natural soluble neuropilin-1 that binds vascular endothelial growth factor: In vivo expression and antitumor activity. Proc. Natl. Acad. Sci. U. S. A. 97, 2573–2578 (2000).

21. Bechara, A. et al. FAK-MAPK-dependent adhesion disassembly downstream of L1 contributes to semaphorin3A-induced collapse. EMBO J. 27, 1549–1562 (2008).

22. Takahashi, T. et al. Plexin-neuropilin-1 complexes form functional semaphorin-3A receptors. Cell 99, 59–69 (1999).

23. Daviaud, N., Chen, K., Huang, Y., Friedel, R. H. & Zou, H. Impaired cortical neurogenesis in plexin-B1 and -B2 double deletion mutant. Dev. Neurobiol. 76, 882–899 (2016).

24. Schwarz, Q. et al. Plexin A3 and plexin A4 convey semaphorin signals during facial nerve development. Dev. Biol. 324, 1–9 (2008).

25. Giger, R. J. et al. Neuropilin-2 is required in vivo for selective axon guidance responses to secreted semaphorins. Neuron 25, 29–41 (2000).

26. Gu, C. et al. Neuropilin-1 conveys semaphorin and VEGF signaling during neural and cardiovascular development. Dev. Cell 5, 45–57 (2003).

27. Shimizu, M., Murakami, Y., Suto, F. & Fujisawa, H. Determination of cell adhesion sites of neuropilin-1. J. Cell Biol. 148, 1283–1293 (2000).

28. Olsson, A.-K., Dimberg, A., Kreuger, J. & Claesson-Welsh, L. VEGF receptor signalling - in control of vascular function. Nat. Rev. Mol. Cell Biol. 7, 359–371 (2006).

29. Casazza, A., Fazzari, P. & Tamagnone, L. Semaphorin signals in cell adhesion and cell migration: functional role and molecular mechanisms. Adv. Exp. Med. Biol. 600, 90–108 (2007).

30. Morgan-Smith, M., Wu, Y., Zhu, X., Pringle, J. & Snider, W. D. GSK-3 signaling in developing cortical neurons is essential for radial migration and dendritic orientation. Elife 3, e02663 (2014).

31. Eickholt, B. J., Walsh, F. S. & Doherty, P. An inactive pool of GSK-3 at the leading edge of growth cones is implicated in Semaphorin 3A signaling. J. Cell Biol. 157, 211–217 (2002).

32. Uchida, Y. et al. Semaphorin3A signalling is mediated via sequential Cdk5 and GSK3beta phosphorylation of CRMP2: implication of common phosphorylating mechanism underlying axon guidance and Alzheimer’s disease. Genes Cells 10, 165–179 (2005).

33. Ng, T. et al. Neuropilin 2 Signaling Is Involved in Cell Positioning of Adult-born Neurons through Glycogen Synthase Kinase-3beta (GSK3beta). J. Biol. Chem. 291, 25088–25095 (2016).

34. Yokota, Y. et al. Cdc42 and Gsk3 modulate the dynamics of radial glial growth, inter-radial glial interactions and polarity in the developing cerebral cortex. Development 137, 4101–4110 (2010).

35. Junghans, D., Hack, I., Frotscher, M., Taylor, V. & Kemler, R. Beta-catenin-mediated cell-adhesion is vital for embryonic forebrain development. Dev. Dyn. an Off. Publ. Am. Assoc. Anat. 233, 528–539 (2005).

36. Hur, E.-M. & Zhou, F.-Q. GSK3 signalling in neural development. Nat. Rev. Neurosci. 11, 539–551 (2010).

37. Arvanitis, D. N. et al. Ephrin B1 maintains apical adhesion of neural progenitors. Development 140, 2082–2092 (2013).

38. Loulier, K. et al. beta1 integrin maintains integrity of the embryonic neocortical stem cell niche. PLoS Biol. 7, e1000176 (2009).

39. Kim, S. et al. Alpha-Synuclein Suppresses Retinoic Acid-Induced Neuronal Differentiation by Targeting the Glycogen Synthase Kinase-3beta/beta-Catenin Signaling Pathway. Mol. Neurobiol. 55, 1607–1619 (2018).

40. Ma, Y.-X. et al. Differential Roles of Glycogen Synthase Kinase 3 Subtypes Alpha and Beta in Cortical Development. Front. Mol. Neurosci. 10, 391 (2017).

41. Bulfone, A. et al. Expression pattern of the Tbr2 (Eomesodermin) gene during mouse and chick brain development. Mech. Dev. 84, 133–138 (1999).

42. Englund, C. et al. Pax6, Tbr2, and Tbr1 are expressed sequentially by radial glia, intermediate progenitor cells, and postmitotic neurons in developing neocortex. J. Neurosci. 25, 247–251 (2005).

43. Hevner, R. F. From radial glia to pyramidal-projection neuron: transcription factor cascades in cerebral cortex development. Mol. Neurobiol. 33, 33–50 (2006).

44. Pontious, A., Kowalczyk, T., Englund, C. & Hevner, R. F. Role of intermediate progenitor cells in cerebral cortex development. Dev. Neurosci. 30, 24–32 (2008).

45. Kriegstein, A. R. Constructing circuits: neurogenesis and migration in the developing neocortex. Epilepsia 46 Suppl 7, 15–21 (2005).

46. Sessa, A., Mao, C.-A., Hadjantonakis, A.-K., Klein, W. H. & Broccoli, V. Tbr2 directs conversion of radial glia into basal precursors and guides neuronal amplification by indirect neurogenesis in the developing neocortex. Neuron 60, 56–69 (2008).

47. Redzic, Z. B., Preston, J. E., Duncan, J. A., Chodobski, A. & Szmydynger-Chodobska, J. The choroid plexus-cerebrospinal fluid system: from development to aging. Curr. Top. Dev. Biol. 71, 1–52 (2005).

48. Zhang, F., Chen, J., Zhao, L. & Dong, C. Candidate biomarkers of multiple system atrophy in cerebrospinal fluid. Rev. Neurosci. 25, 653–662 (2014).

49. Abdi, F. et al. Detection of biomarkers with a multiplex quantitative proteomic platform in cerebrospinal fluid of patients with neurodegenerative disorders. J. Alzheimers. Dis. 9, 293–348 (2006).

50. Chen, H., Chedotal, A., He, Z., Goodman, C. S. & Tessier-Lavigne, M. Neuropilin-2, a novel member of the neuropilin family, is a high affinity receptor for the semaphorins Sema E and Sema IV but not Sema III. Neuron 19, 547–559 (1997).

51. Romi, E. et al. ADAM metalloproteases promote a developmental switch in responsiveness to the axonal repellant Sema3A. Nat. Commun. 5, 4058 (2014).

52. Chauvet, S. et al. Gating of Sema3E/PlexinD1 signaling by neuropilin-1 switches axonal repulsion to attraction during brain development. Neuron 56, 807–822 (2007).

53. Janssen, B. J. C. et al. Neuropilins lock secreted semaphorins onto plexins in a ternary signaling complex. Nat. Struct. Mol. Biol. 19, 1293–1299 (2012).

54. Deng, S. et al. Plexin-B2, but not Plexin-B1, critically modulates neuronal migration and patterning of the developing nervous system in vivo. J. Neurosci. 27, 6333–6347 (2007).

55. Hirschberg, A. et al. Gene deletion mutants reveal a role for semaphorin receptors of the plexin-B family in mechanisms underlying corticogenesis. Mol. Cell. Biol. 30, 764–780 (2010).

56. Fazzari, P. et al. Plexin-B1 plays a redundant role during mouse development and in tumour angiogenesis. BMC Dev. Biol. 7, 55 (2007).

57. Hota, P. K. & Buck, M. Plexin structures are coming: opportunities for multilevel investigations of semaphorin guidance receptors, their cell signaling mechanisms, and functions. Cell. Mol. Life Sci. 69, 3765–3805 (2012).

58. Alto, L. T. & Terman, J. R. Semaphorins and their Signaling Mechanisms. Methods Mol. Biol. 1493, 1–25 (2017).

59. Pascoe, H. G. et al. Secondary PDZ domain-binding site on class B plexins enhances the affinity for PDZ-RhoGEF. Proc. Natl. Acad. Sci. U. S. A. 112, 14852–14857 (2015).

60. Lehtinen, M. K. et al. The cerebrospinal fluid provides a proliferative niche for neural progenitor cells. Neuron 69, 893–905 (2011).

61. Schenk, J., Wilsch-Brauninger, M., Calegari, F. & Huttner, W. B. Myosin II is required for interkinetic nuclear migration of neural progenitors. Proc. Natl. Acad. Sci. U. S. A. 106, 16487–16492 (2009).

62. Clark, B. S. et al. Loss of Llgl1 in retinal neuroepithelia reveals links between apical domain size, Notch activity and neurogenesis. Development 139, 1599–1610 (2012).

63. Baye, L. M. & Link, B. A. Interkinetic nuclear migration and the selection of neurogenic cell divisions during vertebrate retinogenesis. J. Neurosci. 27, 10143–10152 (2007).

64. Del Bene, F., Wehman, A. M., Link, B. A. & Baier, H. Regulation of neurogenesis by interkinetic nuclear migration through an apical-basal notch gradient. Cell 134, 1055–1065 (2008).

65. Latasa, M. J., Cisneros, E. & Frade, J. M. Cell cycle control of Notch signaling and the functional regionalization of the neuroepithelium during vertebrate neurogenesis. Int. J. Dev. Biol. 53, 895–908 (2009).

66. Murciano, A., Zamora, J., Lopez-Sanchez, J. & Frade, J. M. Interkinetic nuclear movement may provide spatial clues to the regulation of neurogenesis. Mol. Cell. Neurosci. 21, 285–300 (2002).

67. Strzyz, P. J. et al. Interkinetic nuclear migration is centrosome independent and ensures apical cell division to maintain tissue integrity. Dev. Cell 32, 203–219 (2015).

68. Alexandre, P., Reugels, A. M., Barker, D., Blanc, E. & Clarke, J. D. W. Neurons derive from the more apical daughter in asymmetric divisions in the zebrafish neural tube. Nat. Neurosci. 13, 673–679 (2010).

69. Mora-Bermudez, F. et al. Differences and similarities between human and chimpanzee neural progenitors during cerebral cortex development. Elife 5, (2016).

70. Pilaz, L.-J. et al. Prolonged mitosis of neural progenitors alters cell fate in the developing brain. Neuron 89, 83–99 (2016).

71. Gallo, G. RhoA-kinase coordinates F-actin organization and myosin II activity during semaphorin-3A-induced axon retraction. J. Cell Sci. 119, 3413–3423 (2006).

72. Takamatsu, H. et al. Semaphorins guide the entry of dendritic cells into the lymphatics by activating myosin II. Nat. Immunol. 11, 594–600 (2010).

73. Foerster, P. et al. mTORC1 signaling and primary cilia are required for brain ventricle morphogenesis. Development 144, 201–210 (2017).

74. Tufro, A. Podocyte Shape Regulation by Semaphorin 3A and MICAL-1. Methods Mol. Biol. 1493, 393–399 (2017).

75. Giacobini, P. et al. Brain endothelial cells control fertility through ovarian-steroid-dependent release of semaphorin 3A. PLoS Biol. 12, e1001808 (2014).

76. Canto, E. et al. Validation of semaphorin 7A and ala-beta-his-dipeptidase as biomarkers associated with the conversion from clinically isolated syndrome to multiple sclerosis. J. Neuroinflammation 11, 181 (2014).

77. Cardenas, A. et al. Evolution of Cortical Neurogenesis in Amniotes Controlled by Robo Signaling Levels. Cell 174, 590-606.e21 (2018).

78. Borrell, V. & Calegari, F. Mechanisms of brain evolution: regulation of neural progenitor cell diversity and cell cycle length. Neurosci. Res. 86, 14–24 (2014).

79. Pilz, G.-A. et al. Amplification of progenitors in the mammalian telencephalon includes a new radial glial cell type. Nat. Commun. 4, 2125 (2013).

